# Transient activation of potent progenitor cells is required for spinal cord regeneration

**DOI:** 10.64898/2026.02.04.703854

**Authors:** Chase A. Weinholtz, Lili Zhou, Vishnu Muraleedharan Saraswathy, Yuxiao Xu, Dana Klatt Shaw, Anthony R. McAdow, Dongkook Park, Jimann Shin, Lila Solnica-Krezel, Aaron N Johnson, Mayssa H. Mokalled

## Abstract

Adult zebrafish exhibit full recovery following spinal cord injury. Transient expansion of stem cell-like progenitors is thought to underlie their regenerative capacity. Yet, our understanding of the identities and contributions of the crucial stem cell populations that direct spontaneous neural repair remains limited. Moreover, while most neural regeneration research is centered on promoting proliferative repair, the regulatory mechanisms that reinstate quiescence post-repair are unknown. Here, we determined the molecular identities and cellular contributions of *sox2*^+^ progenitors during spinal cord repair. Genetic lineage tracing shows zebrafish spinal progenitors, while quiescent in uninjured tissues, self-renew and differentiate into neurons and glia after injury. By single-cell sequencing, *sox2^+^* cells are heterogeneous and biased towards neuronal or glial fates in both homeostatic and regenerating tissues. By screening for transcription factors that are differentially expressed in acute versus chronic spinal cord injury, we find the Bach1 transcription factors control transient progenitor cell activation by acting as dual activators and repressors of *sox2* expression. This study elucidates the molecular diversity and contributions of *sox2* expressing cells during spinal cord repair and identifies a transcriptional regulatory switch by which progenitor cells expand after injury and restore quiescence after regeneration is completed.

## INTRODUCTION

Spinal cord injury (SCI) triggers distinct injury-responsive genetic and cellular programs that underlie differential regenerative capacities across vertebrates (Silver, 2016; Edwards-Faret et al., 2017; Tazaki et al., 2017; Sofroniew, 2018; Maden and Varholick, 2020; Phipps et al., 2020; Walker and Echeverri, 2022; Zheng and Tuszynski, 2023; Tendolkar and Mokalled, 2025). In regenerative vertebrates including zebrafish, spontaneous spinal cord (SC) repair is driven by potent stem cell-like progenitors that surround the central canal and undergo extensive proliferation following injury (Johansson et al., 1999; Mothe and Tator, 2005; Meletis et al., 2008; Barnabe-Heider et al., 2010; Hui et al., 2010; Rodrigo Albors et al., 2023). In non-regenerative mammalian SCs, adult ependymal cells proliferate and differentiate into astrocytes and oligodendrocytes, but their ability to contribute to neuronal differentiation *in vivo* is limited (Martens et al., 2002; Horky et al., 2006; Meletis et al., 2008; Barnabe-Heider et al., 2010; Karimi-Abdolrezaee et al., 2012; McDonough and Martinez-Cerdeno, 2012; Sabelström et al., 2013; Su et al., 2014; Paniagua-Torija et al., 2018; Shah et al., 2018; Xue et al., 2022). Notably, mammalian ependymal cells can be reprogrammed to enhance their potency, suggesting their regenerative potential is transcriptionally dormant (Shihabuddin et al., 2000; Ohori et al., 2006; Lee et al., 2013; Xu et al., 2017; Tai et al., 2021). We propose that elucidating the transcriptional programs that underscore the potency of endogenous stem cells in regenerative animals will guide protocols to improve reprogramming strategies and SCI outcomes.

Injury-induced expression of *sex determining region Y box 2 (sox2)* is a defining feature of neural progenitors in zebrafish (Adolf et al., 2006; Grandel et al., 2006; Kroehne et al., 2011; Barbosa et al., 2015). Sox2 is a highly conserved transcription factor capable of inducing and maintaining pluripotency (Avilion et al., 2003; Graham et al., 2003; Takahashi and Yamanaka, 2006; Schaefer and Lengerke, 2020). In mouse SCI, exogenous *sox2* expression is sufficient to reprogram NG2 glia into neurons *in vivo* (Su et al., 2014). In zebrafish, *sox2*^+^ progenitors elicit compartmentalized responses after SCI (Reimer et al., 2008; Klatt Shaw et al., 2021; Saraswathy et al., 2022; Zhou et al., 2023). For instance, progenitors within the progenitor motor neuron (pMN) domain express *olig2* and regenerate motor neurons (Reimer et al., 2008). A distinct neurogenic niche dorsal to the ependyma upregulates TGF-β signaling to control the rates of self-renewal and interneuron neurogenesis after injury (Saraswathy et al., 2022). On the other hand, ventral *sox2*^+^ progenitors express *connective tissue growth factor a* (*ctgfa*), undergo epithelial-to-mesenchymal transition (EMT) and direct regenerative glial bridging after SCI (Mokalled et al., 2016; Klatt Shaw et al., 2021; Zhou et al., 2023). As these studies revealed niches of *sox2^+^* progenitors underlying the regeneration of distinct cell types following SCI, several fundamental questions remain unresolved: How do SC progenitor niches balance lineage restriction and potency; how many progenitor domains are needed to regenerate spinal cell types; and does the adult zebrafish SC harbor multipotent progenitors? Comprehensive molecular and transcriptional characterization is thus required to determine the potency and contributions of *sox2*^+^ progenitors during innate SC repair.

*sox2* expression and the proliferation of *sox2*^+^ progenitors are transiently activated after SCI. Zebrafish spinal progenitors upregulate *sox2* and proliferate as early as 1 day post-injury (dpi) (Ogai et al., 2014; Hui et al., 2015; Gorsuch et al., 2017). For decades, their acute activation post-injury and their neurogenic potential during sub-acute SCI have been the primary focus of pre-clinical regeneration studies (Becker et al., 2004; Reimer et al., 2013; Barreiro-Iglesias et al., 2015; Hui et al., 2015; Mokalled et al., 2016; Ribeiro et al., 2017; Cavone et al., 2021; Klatt Shaw et al., 2021; Vandestadt et al., 2021; Saraswathy et al., 2022; Cigliola et al., 2023; Zhou et al., 2023; de Sena-Tomas et al., 2024). However, successful SC repair also relies on restricting progenitor cell proliferation and regaining homeostasis as regeneration concludes. Understanding how to re-establish progenitor cell quiescence is also necessary for the development of regenerative therapies. However, the regulatory mechanisms that suppress *sox2* expression and reinstate progenitor cell quiescence as SC repair concludes remain undefined.

BTB and CNC homology 1 (Bach1) is a basic leucine zipper transcription factor that plays important and diverse roles in stem cell pluripotency and differentiation (Itoh-Nakadai et al., 2014; Sato et al., 2020; Suzuki et al., 2020). Bach1 is comprised of several functional domains including an N-terminal BTB/POZ domain that facilitates protein-protein interactions, and a C-terminal basic leucine zipper (bZIP) domain responsible for DNA binding. This structural organization allows Bach1 to form various protein heterodimers that carry out context-dependent transcriptional activation or repression functions (Turner and Tjian, 1989; Albagli et al., 1995; Oyake et al., 1996; Katsani et al., 1999; Sun et al., 2002; Jia et al., 2022; He et al., 2023). Previous studies reported *bach1* expression in ventral spinal progenitors and showed that *bach1a/b* paralogues are required for functional recovery following SCI (Klatt Shaw et al., 2021). However, the timing, transcriptional activity and mechanisms of Bach1 functions during SC repair are still unknown.

This study determines the molecular identities, stem cell properties and contributions of *sox2*^+^ progenitors during SC repair. Using a new *sox2-CreER^T2^* knock-in line for genetic lineage tracing, we show that *sox2*^+^ cells self-renew and give rise to neurons and glia after SCI. Single nuclear RNA sequencing (snRNA-seq) and *in vivo* validation show *sox2^+^*cells are heterogeneous and biased towards distinct neuronal or glial fates in both homeostatic and regenerating SC tissues. To identify transcriptional regulators that play repressive roles in chronic SCI, we screen for transcription factors that are differentially expressed in acute versus chronic SCI. Loss-and gain-of-function studies reveal Bach1a/b as early activators and late repressors of *sox2* expression and progenitor cell expansion. Using a series of luciferase, chromatin immunoprecipitation and genetic rescue assays, we establish Bach1 as a direct transcriptional regulator of *sox2* that de-represses or represses *sox2* expression depending on cofactor availability. These findings delineate the molecular diversity and contributions of *sox2* expressing progenitors during spontaneous SC repair and reveal a regulatory switch can promote or restrain progenitor cell activation at different stages of regeneration.

## RESULTS

### Sox2^+^ cells mount an acute proliferative response in sub-acute SCI

To examine the stem cell properties of Sox2^+^ cells during SC regeneration, wild-type zebrafish adults were subjected to complete SC transections followed by daily intraperitoneal EdU injections after injury. This EdU labeling regimen establishes a cumulative profile of injury-induced cell proliferation. SC tissues at 7, 14 and 28 dpi were compared to tissues collected from uninjured siblings after 6 days of daily EdU injection **(Fig. 1A)**. SC sections 750, 450 and 150 µm rostral and caudal to the lesion were analyzed. Compared to uninjured controls where 2-3% of SC nuclei were EdU^+^, cell proliferation robustly increased after SCI and peaked at 7 dpi (67% EdU incorporation at the lesion) **(Fig. S1A, S1B)**. Mirroring global cell proliferation, the proportion of Sox2^+^ EdU^+^ cells increased from 1-3% in uninjured controls to 72% at 7 dpi, 86% at 14 dpi and began to decline at 28 dpi **(Fig. 1B, 1C)**. These findings suggested Sox2^+^ cells are relatively quiescent in homeostatic adult SCs, and mount an acute and transient proliferative response within 7-14 days of SCI.

**Figure 1.**
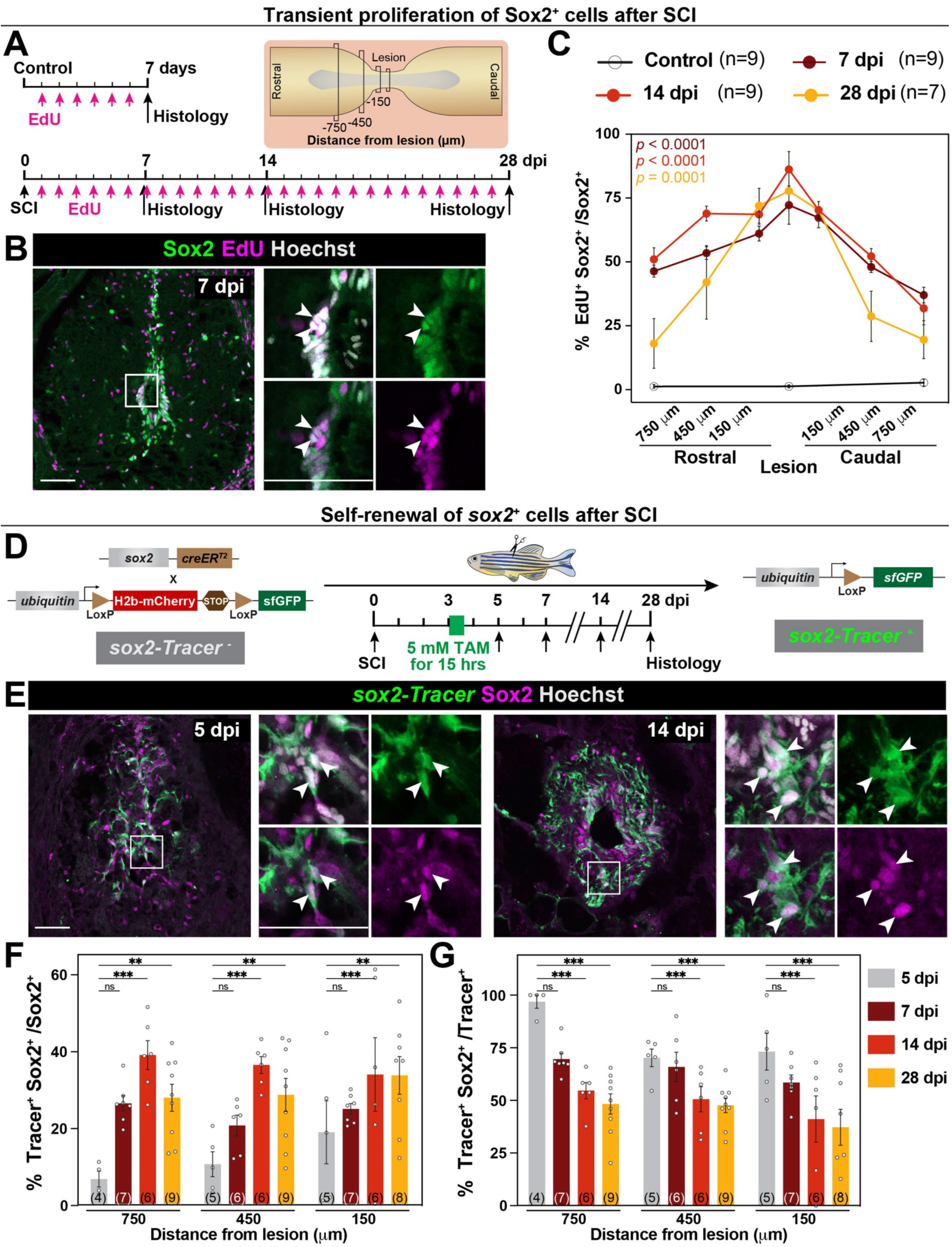
Sox2^+^ cells undergo transient self-renewal after SCI. **(A)** Experimental timeline to evaluate the proliferation of Sox2^+^ cells after SCI. Uninjured SC tissues were collected 6 days after daily EdU injections. Injured tissues were collected at 7, 14 and 28 dpi for histological examination. **(B)** Immunostaining for Sox2 (green), EdU (magenta) and Hoechst (grey) at 7 dpi. SC cross sections 750 µm from the lesion are shown. High-magnification insets show select Sox2^+^ EdU^+^ cells in triple, double or single channel views. Arrowheads indicate Sox2^+^ EdU^+^ cells. **(C)** Quantification of the numbers of Sox2^+^ EdU^+^ cells at 0, 7, 14 and 28 dpi. Percent Sox2^+^ EdU^+^ cells were normalized to Sox2^+^ cells for each section. Cross sections 150, 450 and 750 µm rostral and caudal to the lesion were analyzed. **(D)** Experimental timeline for genetic lineage tracing of Sox2^+^ cells after SCI. *sox2-Tracer* animals were treated with 5 mM Tamoxifen (TAM) for 15 hrs at 3 dpi. SC tissue sections were analyzed at 5, 7, 14 and 28 dpi. **(E)** Immunostaining for *sox2-Tracer*^+^ (GFP, green), Sox2 (magenta) and Hoechst (grey). SC sections 450 µm from the lesion are shown at 5 and 14 dpi. Arrowheads indicate *sox2*-Tracer^+^ Sox2^+^ cells. **(F,G)** Quantification of *sox2-Tracer*^+^ Sox2^+^ cells at 5, 7, 14 and 28 dpi. For each section, the numbers of *sox2-Tracer*^+^ Sox2^+^ cells were normalized to either Sox2^+^ cells (F), or to *sox2-Tracer*^+^ cells (G). Quantifications were performed 150, 450 and 750 µm from the lesion. Data points indicate individual animals and sample sizes are indicated in parentheses. Two-way ANOVA was performed in C. Two-way ANOVA with Holm-Šidák’s multiple comparisons were performed in J and L. Error bars represent SEM. ns, p≥0.05; **p≤0.01; ***p≤0.001. Scale bars: 50 μm, inset 10 μm.

### Sox2^+^ cells self-renew and differentiate following SCI

To determine the potency and contribution of *sox2^+^* cells after SCI, we employed a genetic lineage tracing system that combines a new (*sox2-2a-CreER^T2^)^stl374^* knock-in allele with *ubi:loxP-H2b-mCherry-STOP-loxP-sfGFP^stl345^*, referred to hereafter as *sox2-Tracer* fish (Dean et al., 2025). This system enabled us to induce Tamoxifen-and Cre-dependent recombination to permanently label *sox2*^+^ cells and their progenies during regeneration. To test the specificity of GFP expression after recombination, *sox2-Tracer* fish were subjected to SCI, treated with 5 mM Tamoxifen (TAM) for 15 hours at 3 dpi, and SC tissues were collected at 7 dpi. GFP^+^ cells were only detected in TAM-treated Cre^+^ SCs compared to TAM-treated Cre^-^, vehicle-treated Cre^+^ or vehicle-treated Cre^-^controls **(Fig. S1C, S1D)**. These findings confirmed our TAM treatment induces specific recombination and GFP expression in *sox2^+^* cells.

To determine the potency of *sox2^+^* cells, we performed the same regimen of SCI and TAM treatment at 3 dpi **(Fig. 1D)**. SC tissues were collected at 5, 7, 14 and 28 dpi and stained for *sox2-Tracer* (GFP) and Sox2 **(Fig. 1E, S1E)**. By quantifying *sox2*-derived cells within Sox2^+^ cells (*sox2-Tracer*^+^ Sox2^+^ cells normalized to Sox2^+^ cells), the proportion of recombined cells within Sox2^+^ cells significantly increased from 5 dpi to 14 dpi **(Fig. 1F).** These findings indicated recombined *sox2*-derived cells undergo self-renewal to expand the pool of Sox2^+^ cells after SCI. On the other hand, by quantifying Sox2+ cells within *sox2*-derived cells (*sox2-Tracer*^+^Sox2^+^ cells normalized to *sox2-Tracer*^+^ cells), the proportion of recombined cells that maintained Sox2 expression showed a significant decrease from 5 dpi to 14 and 28 dpi **(Fig. 1G).** These findings indicate a subset of *sox2*-derived cells downregulate Sox2 expression over the regeneration timeline, ostensibly as they differentiate into regenerating cell types. Together, our genetic lineage tracing showed *sox2^+^* cells undergo proliferative self-renewal in sub-acute SCI and downregulate *sox2* expression at later stages of SC regeneration.

### *sox2* is a comprehensive marker of SC progenitors

We performed snRNA-seq to determine the molecular signatures that underlie the potency and transient activation of progenitor cells after SCI. Multiple genes including *sox2, gfap*, *nes* and *foxj1a* have been previously used to label and study zebrafish SC progenitors (März et al., 2010; Kroehne et al., 2011; Ogai et al., 2014; Ribeiro et al., 2017; Than-Trong et al., 2018; Than-Trong et al., 2020). Postulating that *sox2* serves as one of the most comprehensive markers of SC progenitors, we performed snRNA-seq using an established *sox2-2a-sfGFP^stl84^*knock-in line (Shin et al., 2014). Despite the technical challenges of co-localizing cytoplasmic *sox2-2a-sfGFP* expression with nuclear Sox2 protein, we confirmed that >80% Sox2^+^ nuclei were sfGFP^+^ and vice versa **(Fig. 2A)**. For snRNA-seq, *sox2-2a-sfGFP* fish were subjected to SCI and SC tissues spanning 3 mm around the lesion site were collected at 7 and 14 dpi **(Fig. 2B)**. Comparable SC tissue sections were collected from uninjured controls. For each time point, 2 replicate pools of nuclei were sequenced, and 50 fish were pooled within each replicate. A total of 63,804 nuclei were sequenced **(Fig. 2B)**. After quality control analysis **(Fig. S2A, S2B)**, unsupervised Seurat clustering yielded 33 clusters including neurons, glia, oligodendrocytes and OPCs **(Fig. 2C)**. Gene expression analysis of various progenitor markers (*sox2, gfap, foxj1a, cspg4, prom2* and *nes*) in this integrated dataset showed pronounced differences that reflect progenitor cell diversity in SC tissues **(Fig. 2D, S2C)**. For instance, *cspg4*, which is expressed in bipotent OPCs that give rise to neurons and oligodendrocytes during embryonic development, and *foxj1a*, which labels ependymal progenitors were expressed in distinct clusters by snRNA-seq **(Fig. S2C)** (Ribeiro et al., 2017; Bromley-Coolidge et al., 2024). Compared to and encompassing other progenitor markers, *sox2* was comprehensively expressed in *nes*^+^ neural stem cells, *cspg4*^+^ OPCs and *foxj1a^+^* ependymal cells **(Fig. 2D, S2C)**. *sox2* was also strongly upregulated across clusters after injury **(Fig. 2E)**. Aiming to broadly capture and characterize SC progenitors, we subclustered *sox2-2a-sfGFP* expressing cells for subsequent analysis.

**Figure 2.**
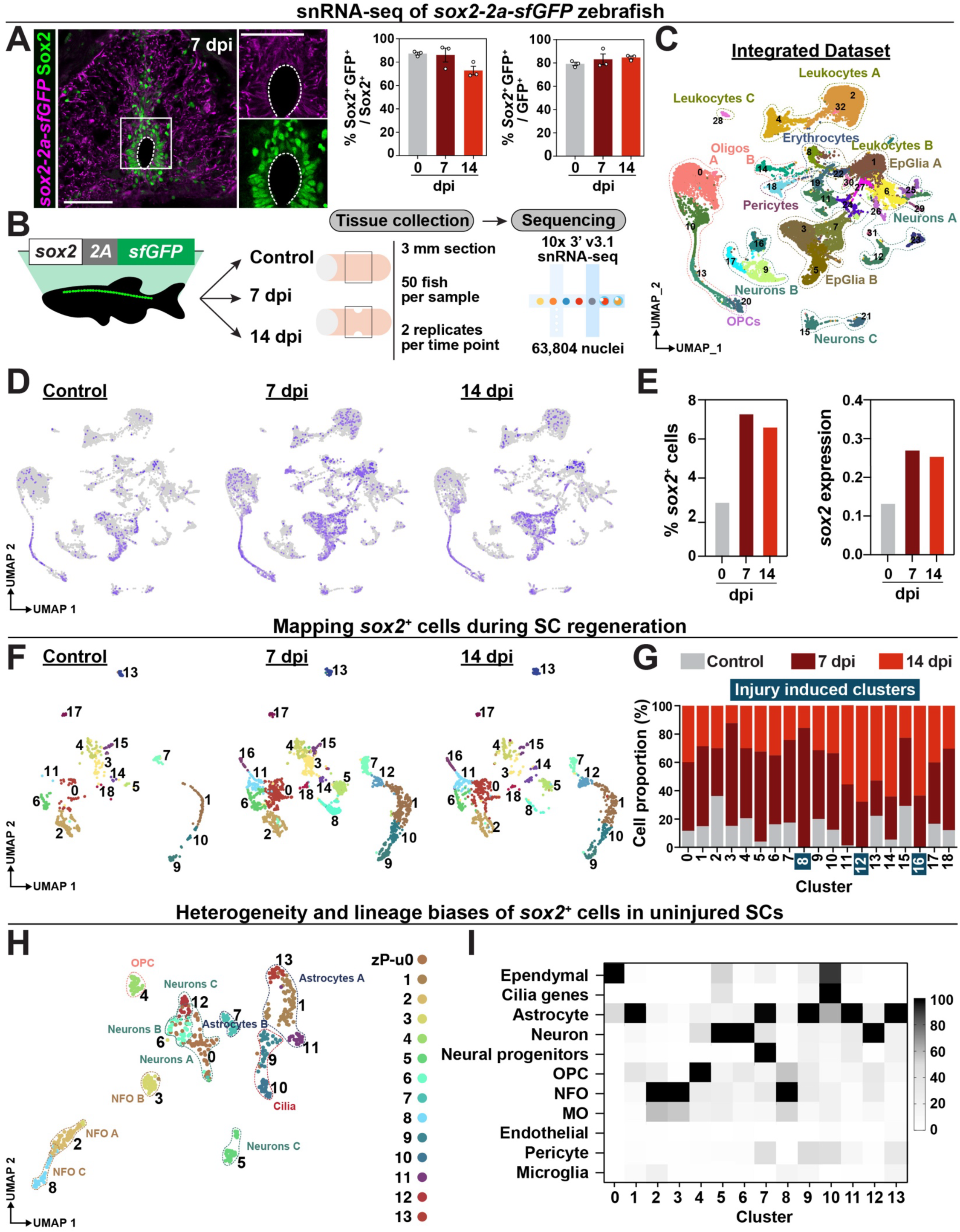
*sox2^+^*cells are heterogeneous and lineage-biased in homeostatic and lesioned SCs. **(A)** Sox2 staining in *sox2-2a-sfGFP* zebrafish at 7 dpi. SC cross sections 450 μm from the lesion are shown and dotted lines delineate central canal edges. The numbers of GFP^+^ Sox2^+^ nuclei were normalized to either Sox2^+^ (left) or GFP^+^ (right) nuclei. **(B)** Experimental pipeline to generate a single-cell atlas of zebrafish SC progenitors. SC tissue collection, nuclear isolation and single nuclear RNA-seq (snRNA-seq) were performed on *sox2-2a-sfGFP* knock-in fish at 0, 7 and 14 dpi. **(C)** Merged UMAP representation of the integrated dataset. Two biological replicates, 3 time points and 63,804 nuclei were clustered into major spinal cell populations. **(D,E)** Feature plots of *sox2^+^* cells during SC regeneration. For each time point, cell proportions were normalized to the total number of cells at that time point. **(F)** Split UMAP representation shows 19 subclusters of *sox2*^+^ cells at 0.6 resolution parameter. Two biological replicates and 3,620 *sox2*^+^ cells were analyzed. **(G)** Distribution of *sox2*^+^ cells per cluster during SC regeneration. For each time point and cluster, the numbers of *sox2*^+^ cells were normalized to the total number of cells within that cluster. **(H)** UMAP representation of *sox2*^+^ cells in uninjured SC tissues. Cluster identities and numbers were matched with their corresponding clusters from the integrated dataset in F. **(I)** Lineage biases of *sox2*^+^ cells in uninjured SC tissues. The top DE markers for each cluster were cross-referenced with our assembled database of vertebrate nervous system markers (VNM marker database). Heatmap depicting-log10 p-value for the marker scoring output from the “DEMarkerScoring” algorithm is shown. One-tailed hypergeometric probability test was used to calculate the p-value.-log10 p-value for each cluster was scaled between 0-100. Cluster identity was assigned based on the highest-log10 p-value obtained for each cluster. Scale bars: 50 μm.

### *sox2*^+^ cells are heterogeneous and lineage-biased before and after SCI

Unsupervised Seurat clustering of *sox2-2a-sfGFP* expressing cells yielded 19 clusters highlighting the transcriptional diversity of *sox2*^+^ cells **(Fig. 2F, S2D-F)**. By assessing changes in cell proportions within each cluster between 0 and 14 dpi, only 3 clusters (8, 12 and 16) were classified to be injury-induced, suggesting they may have regenerative functions during SC repair **(Fig. 2F, 2G)**. Surprisingly, 16 out of 19 *sox2*^+^ cell clusters were present in both injured and uninjured SC, suggesting *sox2*^+^ cells are heterogeneous and lineage-biased in adult zebrafish regardless of injury status. To test the heterogeneity of *sox2*^+^ cells in homeostatic adults, we performed unsupervised Seurat clustering for *sox2-2a-sfGFP* expressing cells in uninjured animals without integrating post-injury datasets. This analysis revealed 14 clusters comprised of diverse cell types, mirroring the cluster identities obtained by integrating uninjured and injured time points **(Fig. 2H)**. These naïve *sox2*^+^ cells displayed differential gene expression congruent with lineage biases towards various glial and neuronal cell types **(Fig. 2I, S2G).** As the SC regeneration field has been focused on studying genes that are induced in different progenitor domains after SCI, the underlying assumption has been that niches of specialized progenitors emerge after SCI and that quiescent progenitors represent a more homogenous transcriptional profile prior to injury. Contradicting this model, our unbiased snRNA-seq indicates naïve *sox2^+^*cells are heterogenous and transcriptionally biased towards specific lineages in homeostatic adult zebrafish.

We performed Hybridization Chain Reaction (HCR) *in situ* hybridization to validate select progenitor cell clusters identified *in silico*. By snRNA-seq, cluster 0 exhibited elevated expression of cilia-related genes, suggesting it is likely to represent ependymal cells **(Fig. S2E)**. *wnt4a* was identified as a unique marker of c0 and plays an important role in cellular differentiation in adult SCs **(Fig. 3A)** (Gao et al., 2016; Rodrigo Albors et al., 2023). Indeed, HCR labeling showed elevated *wnt4a* expression in the ventral and lateral ependymal cells that encircle the central canal **(Fig. 3B)**. *wnt4a* transcripts co-localized *sox2-2a-sfGFP* and were upregulated proximal and rostral to the lesion at 7 and 14 dpi **(Fig. 3C)**. Further validation showed that *pcdh18b*, predicted to label ependymal cluster 11 **(Fig. S2E)**, was also expressed in a subset of central canal lining cells at 7 dpi **(Fig. S3A)**. On the other hand, the transcription factor *lhx4*, responsible for V2a interneuron differentiation in mice, was identified as a unique marker of neuron-biased c16 **(Fig. 3D)** (Renaux et al., 2024). By HCR *in situ* hybridization, *lhx4* expression was detected in domains consistent with neuronal progenitors and co-localized with both *sox2* and the post-mitotic neuronal marker HuC/D **(Fig. 3E-G)**. Further validation mapped neurogenic clusters 4, 13, 15, 16 and 17 to distinct neurogenic domains **(Fig. S3B).** Together, these scRNA-seq and HCR gene expression studies mapped ependymal and neurogenic progenitors within zebrafish SC.

**Figure 3.**
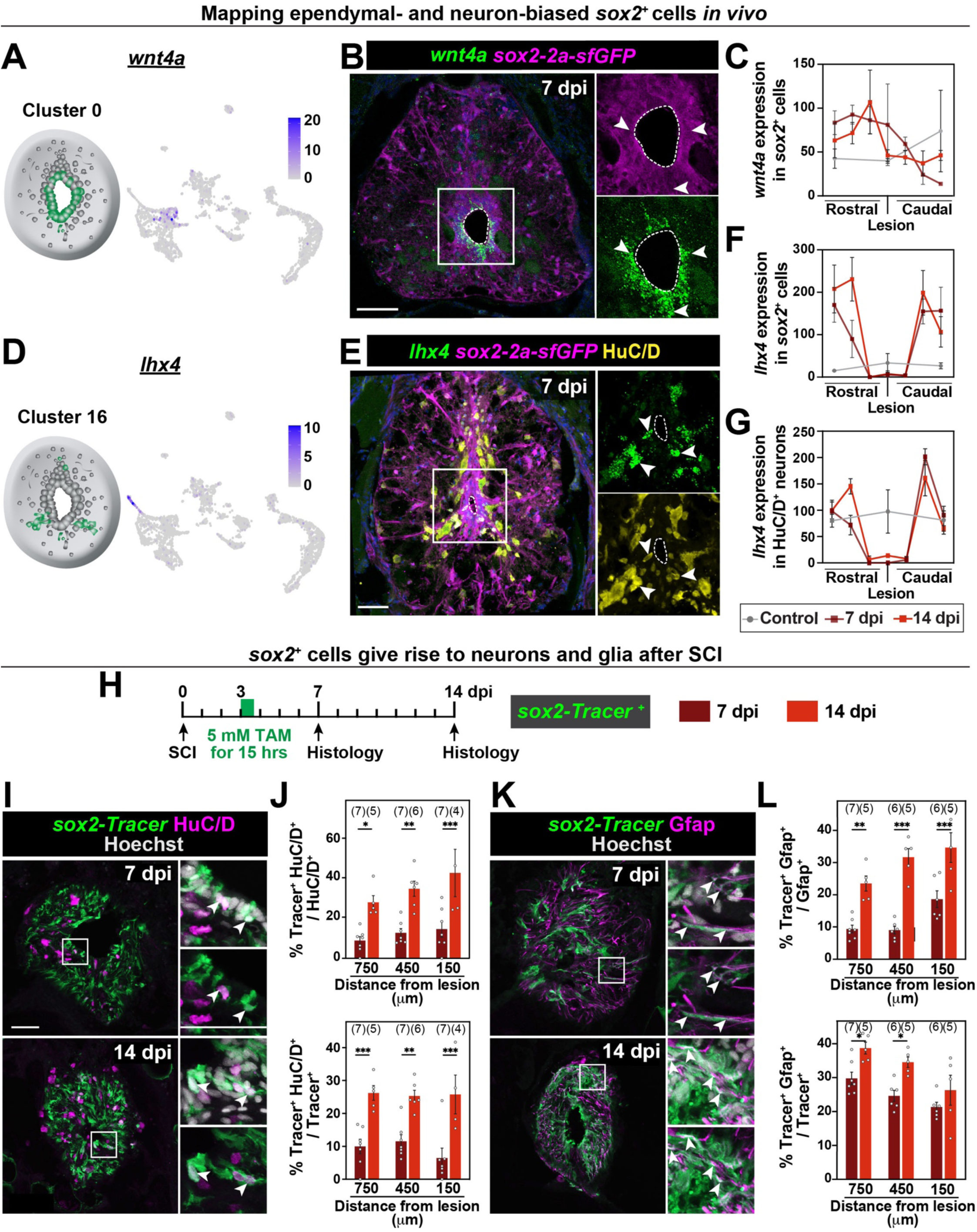
Compartmentalized niches of *sox2*^+^ cells give rise to neurons and glia after SCI. (A-C) *in vivo* validation of ependymal-biased *sox2*^+^ cell cluster 0. Feature plot shows *wnt4a* expression in c0. HCR *in situ* hybridization shows *wnt4a* transcripts (green) at 7 dpi. Arrowheads point to *wnt4a^+^sox2-2a-sfGFP^+^* cells. Colocalization of *wnt4a* with *sox2*-*2a*-*sfGFP* was quantified at 0, 7 and 14 dpi. **(D-G)** *in vivo* validation of neuron-biased *sox2*^+^ cell cluster 16. Feature plot shows *lhx4* expression in c16. HCR *in situ* hybridization shows *lhx4* transcripts (green) at 7 dpi. Arrowheads point to *lhx4^+^sox2-2a-sfGFP^+^*cells. Colocalization of *wnt4a* with *sox2*-*2a*-*sfGFP* (F) and with HuC/D (G) were quantified at 0, 7 and 14 dpi. **(H)** Experimental timeline for genetic lineage tracing of Sox2^+^ cells after SCI. *sox2-Tracer* animals were treated with 5 mM Tamoxifen (TAM) for 15 hrs at 3 dpi. SC tissue sections were analyzed at 7 and 14 dpi. **(I)** Immunostaining for *sox2*-Tracer^+^ (GFP, green), HuC/D (magenta) and Hoechst (grey). SC sections from TAM-treated *sox2*-Tracer (Cre^+^) animals are shown. Insets show high magnification *sox2*-Tracer^+^ cells marked with white arrowheads. **(J)** Quantification of *sox2*-Tracer^+^ and HuC/D^+^ colocalization at 7 and 14 dpi. The numbers of *sox2*-Tracer^+^ and Gfap^+^ cells were quantified. The numbers of s*ox2*-Tracer^+^ HuC/D^+^ cells were normalized to the number of HuC/D^+^ cells (upper panel), and to the total number of *sox2*-Tracer^+^ cells (lower panel). **(K)** Immunostaining for *sox2*-Tracer^+^ (GFP, green), Gfap (magenta) and Hoechst (grey). SC sections from TAM-treated *sox2*-Tracer (Cre^+^) animals are shown. Insets show high magnification *sox2*-Tracer^+^ cells with white arrowheads indicating *sox2*-Tracer^+^ Gfap^+^ staining. **(L)** Quantification of *sox2*-Tracer^+^ and Gfap^+^ colocalization at 7 and 14 dpi. *sox2*-Tracer^+^ and Gfap^+^ fluorescence area was quantified. s*ox2*-Tracer^+^ HuC/D^+^ expression was normalized to Gfap expression (upper panel), and to *sox2*-Tracer expression (lower panel). Data points indicate individual animals and sample sizes are indicated in parentheses. Two-way ANOVA with Holm-Šidák’s multiple comparisons were performed in J and L. Error bars represent SEM. ns, p≥0.05; *p<0.05; **p<0.01; ***p<0.001. Scale bars: 50 μm, inset 10 μm.

### Sox2^+^ progenitors give rise to neurons and glia during SC repair

We used *sox2-CreER^T2^*-based genetic lineage tracing to track the fates of *sox2*^+^ progenitors and their progenies during SC regeneration **(Fig. 3H)**. GFP labeling was co-stained with neuronal (HuC/D) and pan-glial (Gfap) markers **(Fig. 3I-L)**. The proportions of *Tracer^+^* neurons (GFP^+^ HuC/D^+^) increased 3-fold between 7 and 14 dpi, with progenitor-derived neurons comprising 27-42% of all neurons 750 µm from the lesion **(Fig. 3J)**. Consistent with ongoing neurogenesis of *sox2*-derived cells, HuC/D expression within recombined *Tracer^+^* cells increased 3-fold from 7 to 14 dpi **(Fig. 3J)**. Similarly, the proportions of *Tracer^+^* glia (GFP^+^ Gfap^+^) increased 2.6-fold between 7 and 14 dpi **(Fig. 3L).** Progenitor-derived glia comprised 23-34% of all glia 750 µm from the lesion and accounted for 26-38% of total traced cells at 14 dpi **(Fig. 3L)**. Together, our *in silico* and *in vivo* studies underscore a stepwise process whereby lineage-biased quiescent progenitors transiently upregulate *sox2* and self-renew between 0 and 14 dpi, begin to differentiate into neurons and glia starting at 7 dpi, and regain quiescence at later stages of regeneration.

### *bach1a/b* are transcriptionally enriched in *sox2*^+^ cells after SCI

While progenitor cell activation has been widely studied, we aimed to examine how progenitor cells return to quiescence during later regeneration stages. Postulating *sox2* regulation is a major tipping point between quiescence and activation **(Fig. 1)**, we set out to identify putative *sox2* activators and repressors **(Fig. 4A)**. To this end, we performed pseudobulk analysis to identify transcription factors that are enriched at 7 versus 42 dpi **(Fig. 4B)**. Pseudobulk analysis enhanced our ability to capture transcription factors, which exert biologically significant effects but elicit low gene expression levels in single-cell datasets (Pokhilko et al., 2021). This analysis identified 21 transcription factors enriched at 7 dpi, coinciding with peak *sox2* expression. Compared to 7 dpi, 16 transcription factors were identified at 42 dpi, when *sox2* expression returns to pre-injury levels and regeneration concludes. Intriguingly, while activators account for 71% of sub-acutely expressed transcription factors at 7 dpi (15 out of 21), they decrease to 56% of transcription factors expressed at 42 dpi (6 out of 16) **(Fig. 4B).** Notably, we identified *bach1b* as the only dual-function transcription factor that has canonical activator and repressor activity and whose transcripts were detected at both 7 and 42 dpi **(Fig. 4C)**. This analysis identified activators and repressors of gene transcription during sub-acute and chronic SCI, and suggested Bach1 may play dual roles in the initiation and termination of SC regeneration.

**Figure 4.**
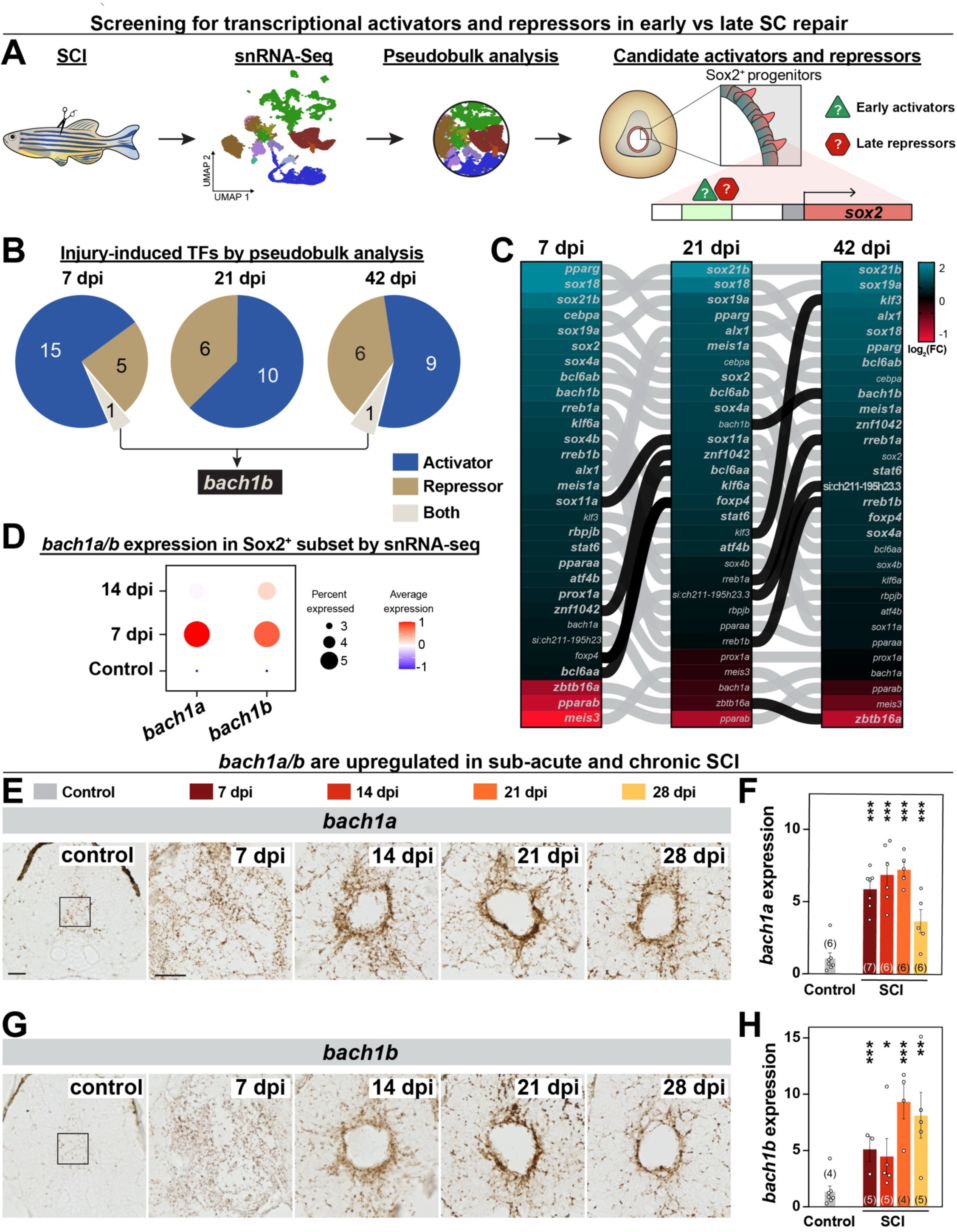
Screening for activators and repressors during SC regeneration. **(A)** Experimental pipeline to identify transcriptional activators and repressors in sub-acute and chronic SCI. Pseudobulk analysis was performed to overcome the limitation of detecting transcription factors by snRNA-seq. **(B)** Proportions of activators, repressors or dual-function transcription factors identified from pseudobulk analysis at 7, 21 and 42 dpi compared to uninjured controls. **(C)** Heatmap depicting the expression of differentially expressed transcription factors at 7, 21 and 42 dpi. Transcription factors that are differentially expressed compared to uninjured at any given time point are bolded. Dark connector lines indicate genes that gained significance between time points. **(D)** Dotplots represent *bach1a* and *bach1b* expression in subclustered *sox2^+^* cells at 0, 7 and 14 dpi. **(E)** RNAscope *in situ* hybridization for *bach1a* during SC regeneration. Cross sections from wild type zebrafish at 0, 7, 14, 21 and 28 dpi are shown. **(F)** Quantification of *bach1a* expression during SC regeneration. *bach1a^+^* area was quantified and normalized to uninjured controls. Sections 450 µm from the lesion were quantified. **(G)** RNAscope *in situ* hybridization for *bach1b* during SC regeneration. Cross sections from wild type zebrafish at 0, 7, 14, 21 and 28 dpi are shown. **(H)** Quantification of *bach1b* expression during SC regeneration. *bach1b^+^*area was quantified and normalized to uninjured controls. Sections 450 µm from the lesion were quantified. Data points indicate individual animals and sample sizes are indicated in parentheses. Two-way ANOVA with Holm-Šidák’s multiple comparisons were performed in F and H. Error bars represent SEM. *p<0.05; **p<0.01; ***p<0.001. Scale bars: 50 μm, inset 10 μm.

By snRNA-seq, *bach1b* and its paralogue *bach1a* were broadly expressed in multiple *sox2^+^* subclusters **(Fig. S4A, S4B)**. Consistent with sustained *bach1b* expression at 7 and 42 dpi, *bach1b* was upregulated at 7 and 14 dpi in pooled subclusters of *sox2^+^*cells **(Fig. 4D)**. On the other hand, *bach1a* was only transiently upregulated in *sox2^+^* cells at 7 dpi **(Fig. 4D)**. *in situ* hybridization confirmed *bach1a* and *bach1b* were markedly upregulated at 7 dpi compared to uninjured controls **(Fig. 4E-H, S4C, S4D)**. *bach1a* maintained elevated expression between 7 and 21 dpi and was downregulated at 28 dpi **(Fig. 4E, 4F)**. Compared to uninjured controls, *bach1b* showed a 5-fold increase at 7 and 14 dpi and a further increase to 9-and 8-fold at 21 and 28 dpi **(Fig. 4G, 4H)**. Notably, *bach1a* and *bach1b* were strongly induced around the central canal, in a pattern consistent with *sox2^+^* progenitors. HCR *in situ* hybridization for *bach1*a/b in *sox2-2A-sfGFP* zebrafish confirmed *bach1a/b* are preferentially upregulated in *sox2^+^* cells, **(Fig. S4E, S4F)**. These findings identified *bach1a/b as* putative regulators of progenitor cell dynamics after SCI.

### *bach1a/b* are required for functional and anatomical regeneration after SCI

To examine the role of *bach1a*/b during functional recovery, we performed SC transections on *bach1a^stl666^;bach1b^stl668^*double mutant zebrafish, referred to hereafter as *bach1a/b* mutants or *bach1a/b^-/-^* (Klatt Shaw and Mokalled, 2021). We then evaluated functional recovery at 14, 28 and 42 dpi **(Fig. 5A)(Mokalled et al., 2016)**. Siblings that are double heterozygous for *bach1a* and *bach1b* (*bach1a/b^+/-^*) were used as controls. Swim endurance was comparable across cohorts of uninjured fish and declined in both groups at 14 dpi. *bach1a/b^+/-^* animals regained swim function at 28 and 42 dpi. In contrast, *bach1a/b* mutants did not recover any swim function after 14 dpi and showed significantly lower swim endurance compared to controls at 28 and 42 dpi **(Fig. 5B)**. To assess anatomical regeneration, rostral neurons were anterogradely traced with biocytin and visualized caudal to the lesion (Klatt Shaw et al., 2021). At 42 dpi, the proportion of caudally traced axons proximal and distal to the lesion was significantly reduced in *bach1a/b^-/-^* compared to *bach1a/b^+/-^*siblings **(Fig. 5C)**. By Gfap staining, glial bridging was deficient in *bach1a/b* mutants relative to controls **(Fig. 5D)** (Klatt Shaw et al., 2021). These results confirmed *bach1a/b* are required for functional and anatomical recovery after SCI.

**Figure 5.**
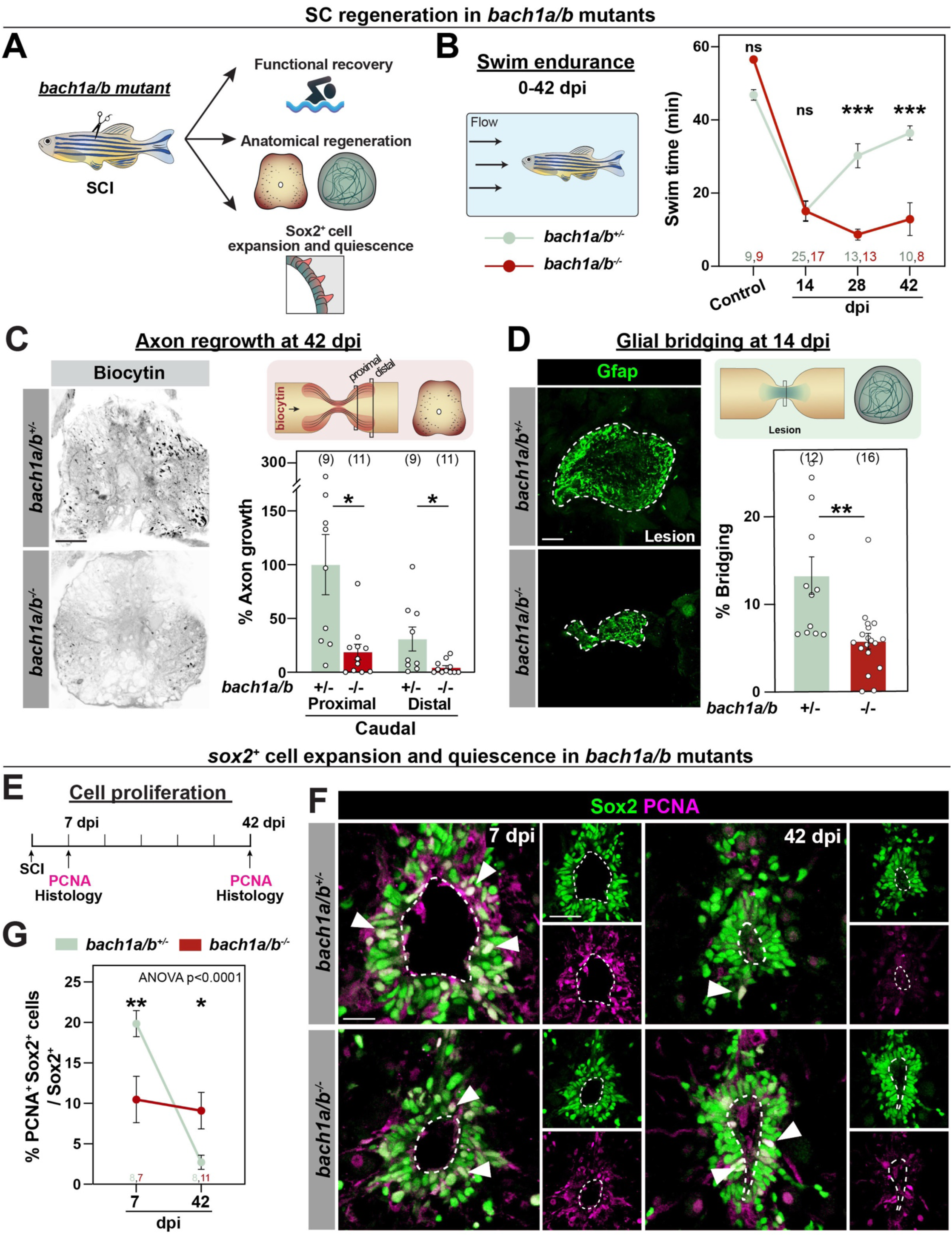
Loss of *bach1* impairs the expansion and quiescence of *sox2*^+^ cells after SCI. **(A)** *bach1a/b* double mutant fish were subjected to SCI. Standard regeneration assays and the profiles of *sox2*^+^ cells were assessed at 7 and 42 dpi. **(B)** Swim endurance for *bach1a/b^-/-^* fish and *bach1a/b^+/-^* control siblings at 14, 28 and 42 dpi. This behavioral test assessed the capacity of *bach1a/b^-/-^* animals to swim in an enclosed swim tunnel under increasing water current velocities. The time at exhaustion for each group is shown. **(C)** Anterograde axon tracing in *bach1a/b^-/-^* and *bach1a/b^+/-^* controls at 42 dpi. Biocytin axon tracer was applied rostrally and analyzed at 600 μm (proximal) and 1500 μm (distal) caudal to the lesion. Representative images show the extent of Biocytin labeling 600 μm (proximal) caudal to the lesion. Axon growth was first normalized to the extent of Biocytin rostral to the lesion, and then to labeling in *bach1a/b^+/-^*controls. **(D)** Glial bridging in *bach1a/b^-/-^* fish and *bach1a/b^+/-^* control siblings at 14 dpi. Representative immunohistochemistry shows the Gfap^+^ bridge at the lesion site. Percent bridging represents the cross-sectional area of the glial bridge at the lesion site relative to the intact SC 750 μm rostral to the lesion. **(E)** Experimental timeline to assess Sox2^+^ cell proliferation at 7 and 42 dpi. **(F)** PCNA (magenta) and Sox2 (green) staining in *bach1a/b^-/-^* and *bach1a/b^+/-^* controls. Representative micrographs 450 µm from the lesion are shown. Arrowheads indicate Sox2^+^ PCNA^+^ cells. **(G)** Quantification of Sox2^+^ cell proliferation at 7 and 42 dpi. For each section, the numbers of PCNA^+^ Sox2^+^ cells were quantified and normalized to Sox2^+^ cells. Quantifications were performed 450 µm from the lesion. For all quantifications, data points indicate individual animals and sample sizes are indicated in parentheses. Two-way ANOVA with Holm-Šidák’s multiple comparisons were performed in B, C and G. Unpaired t test was performed in D. Error bars represent SEM. ns, p≥0.05; *p<0.05; **p<0.01; ***p<0.001. Scale bars: 50 μm.

### *bach1a/b* are required for activation and quiescence of Sox2^+^ progenitors

To examine the role of *bach1a/b* in progenitor activation, we assayed the proliferation of Sox2^+^ cells in *bach1a/b* mutants **(Fig. 5E)**. Sox2 and PCNA staining were performed at 7 and 42 dpi, providing snapshots of actively dividing cells at the time of collection **(Fig. 5F)**. Sox2^+^ PCNA^+^ cells, which accounted for 20% of Sox2^+^ cells in SC sections of *bach1a/b^+/-^* controls at 7 dpi, were 50% depleted in *bach1a/b* mutants **(Fig. 5G, S5A, S5B)**. Consistent with decreased progenitor cell proliferation in sub-acute SCI, daily EdU injections showed the cumulative proportions of Sox2^+^ EdU^+^ cells decreased by 71% in *bach1a/b* mutants at 7 dpi **(Fig. S5C-G)**. Intriguingly, loss of *bach1a/b* caused an inverted phenotype at 42 dpi, whereby the proportions of proliferating Sox2^+^ PCNA^+^ cells increased 3.7-fold in *bach1a/b* mutants compared to controls **(Fig. 5F-G)**. One possibility is that SC regeneration is slower or delayed in the absence of *bach1a/b*, accounting for an extended timeline of Sox2^+^ cell proliferation in *bach1a/b* mutants. Alternatively, as Bach1 can act as a transcriptional repressor and is expressed in chronic SCI, it is plausible that Bach1 is required for both early induction and late repression of Sox2^+^ cell proliferation. To determine whether *sox2* expression is transcriptionally impacted in *bach1a/b* mutants, we performed RT-qPCR for *sox2* on *bach1a/b* mutants at 7 and 42 dpi **(Fig. S5H)**. Mirroring our quantifications of Sox2^+^ cells **(Fig. 5E-G)**, loss of *bach1a/b* depleted *sox2* transcripts at 7 dpi, but significantly enhanced *sox2* expression at 42 dpi **(Fig. S5H)**. These results indicated *bach1a/b* are required to induce Sox2^+^ cells in sub-acute SCI, and to restore their homeostatic quiescence at later stages of SC repair.

### Bach1a/b are direct transcriptional regulators of *sox2*

To examine whether Bach1 is a direct transcriptional regulator of *sox2*, we searched for the consensus binding motif of Bach1 (TGACTCAGC) within a 5 kb region upstream of the *sox2* start codon (Fornes et al., 2020). This analysis identified 8 putative binding regions with a tolerance of 1 nucleotide mismatch **(Fig. 6A)**. We used chromatin immunoprecipitation (ChIP) to assess Bach1 binding at 7 and 42 dpi **(Fig. 6B)**. In the absence of a reliable antibody targeting zebrafish Bach1a/b, we generated a new *hsp70:hBACH1-2A-EGFP^stl692^* transgenic line expressing human BACH1 under control of an inducible heat shock promoter, referred to hereafter as *hBACH1^OE^* fish **(Fig. S6A)**. Expression of hBACH1 and antibody specificity were confirmed by western blot **(Fig. S6B)**. Noting that the *hBACH1^OE^* line did not induce excessive levels of BACH1 overexpression, we proceeded with ChIP. *hBACH1^OE^* fish were subjected to SCI and 3 daily heat shocks preceding collection of SC tissues at either 7 or 42 dpi **(Fig. 6B)**. ChIP-qPCR revealed robust enrichment of hBACH1 at 3 sites (Binding elements 1, 5 and 6) upstream of *sox2* at 7 dpi **(Fig. 6C)**. Interestingly, 1 out the 3 sites (Binding element 5) also enriched for hBACH1 binding at 42 dpi **(Fig. 6C)**. These results showed Bach1 binds distinct regulatory elements upstream of *sox2* at different stages of SC regeneration, and suggested Bach1 regulates *sox2* transcription in sub-acute and chronic SCI.

**Figure 6.**
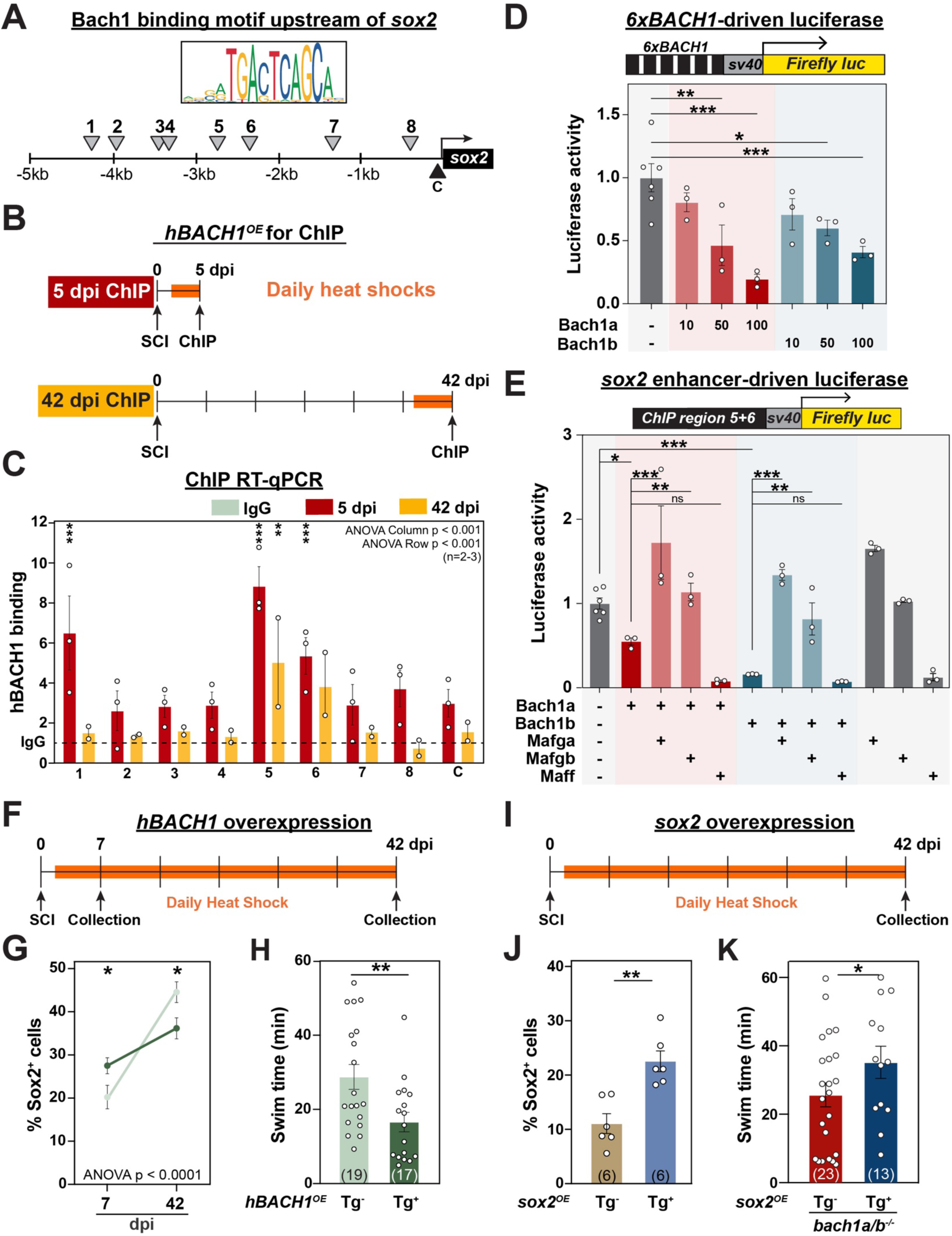
Bach1 is a dual-regulator of *sox2* expression in sub-acute and chronic SCI. **(A**) Schematic representation of the genomic sequence upstream of *sox2*. Grey arrowheads indicate putative binding sites for hBACH1 +/-1 mismatch. The control region marked with black arrowhead was used to detect background binding by ChIP. **(B)** Experimental timeline for ChIP depicting 3 daily heat shocks and SC tissue collection at either 5 or 42 dpi. **(C)** qRT-PCR analysis of hBACH1 enrichment at genomic regions corresponding to those marked in A at 5 and 42 dpi. Y axis represents hBach1 binding. Numbers on the X axis represent putative Bach1 binding sites from 6A, C represents a control genomic region that does not contain Bach1 binding sites. hBach1 binding was calculated using the ΔCq method and normalized to IgG immunoprecipitation (dotted line). **(D)** Luciferase assays using *6xBACH1*-driven Firelfy luciferase. Quantification of *6xBACH1*-Firefly luciferase after co-expression of Bach1a or Bach1b. Y axis represents that activity of Firefly luciferase normalized to control Renilla luciferase and to control wells transfected with empty pcDNA3 vector. **(E)** Luciferase assays using *sox2* enhancer-driven Firelfy luciferase. A 402 bp genomic region containing Bach1 binding regions 5 and 6 upstream of *sox2* was used. Bach1a, Bach1b with or without Maf transcription factors were co-expressed. Firefly luciferase activity was normalized to Renilla luciferase and to control wells transfected with empty pcDNA3 vector. **(F-H)** Experimental timeline for *hBACH1* overexpression. Sox2^+^ cells were quantified 450 µm from the lesion at 7 and 42 dpi (G). For each section, the numbers of Sox2^+^ cells were normalized to the total number of nuclei. Swim endurance for *hBACH1^OE^* fish and control siblings was performed at 42 dpi (H). **(I-K)** Experimental timeline for *sox2* overexpression in *bach1a/b* mutants. Sox2^+^ cells were quantified 450 µm from the lesion (J). Swim endurance for *sox2^OE^;bach1a/b^-/-^*fish and *bach1a/b^-/-^* controls was performed at 42 dpi (K). Data points indicate individual animals or separate biological replicates, and sample sizes are indicated in parentheses. Two-way ANOVA with Holm-Šidák’s multiple comparisons were performed in C and G. Ordinary one-way ANOVA with Holm-Šidák’s multiple comparisons was performed in D and E. Unpaired t tests were performed in J and H. Error bars represent SEM. ns, p≥0.05; *p<0.05; **p<0.01; ***p<0.001.

### Bach1a/b exhibit dual activation and repression activity

We performed a series of luciferase assays to examine whether Bach1 activates and/or represses *sox2* expression after SCI. First, we characterized the inherent transcriptional activity of zebrafish Bach1a and Bach1b using a synthetic enhancer that contains 6 tandem copies of the BACH1 consensus motif (TGACTCAGC) (Fornes et al., 2020). To this end, *6xBACH1* was cloned into the *sv40* minimum promoter-driven luciferase construct and co-transfected into HEK-293T cells with constructs encoding Bach1a or Bach1b. In this experiment, Bach1a and Bach1b repressed luciferase activity in a dose dependent manner **(Fig. 6D)**. In a second set of luciferase assays, a 402 bp region encompassing binding elements 5 and 6 was cloned into the *sv40* minimum promoter-driven luciferase construct, and co-transfected with Bach1a and Bach1b into HEK-293T cells **(Fig. 6E)**. Recapitulating Bach1 repressive activity in the *6xBACH1* luciferase assay, either Bach1a or Bach1b alone significantly repressed baseline luciferase activity **(Fig. 6E)**. As Bach1 is known to partner with small MAF proteins and numerous other cofactor complexes to regulate gene expression (Oyake et al., 1996), we tested whether the transcriptional activity of Bach1 can be modulated by co-expressing zebrafish Mafga, Mafgb or Maff **(Fig. 6E)**. In these assays, Mafga and Mafgb were sufficient to overcome the repressive effects of Bach1a and Bach1b. On the other hand, co-expression with Maff exacerbated Bach1-mediated repression. These findings established context-dependent regulation of *sox2* expression by Bach1a/b and their Maf binding partners.

### BACH1 overexpression alters the expansion and quiescence of *sox2^+^* cells after SCI

Our loss-of-function and molecular studies support a model in which Bach1 performs context-dependent regulatory roles after SCI, activating *sox2* expression in sub-acute SCI and repressing *sox2* at late stages of regeneration. One prediction of this model is that Bach1 overexpression can either enhance or repress progenitor cell activation at different SCI stages. To test this prediction, we performed SCI and daily heat shocks on *hBACH1^OE^* and wild-type siblings **(Fig. 6F)**. We then evaluated the profiles of Sox2^+^ cells at 7 and 42 dpi. Supporting a dual regulatory role for Bach1, *hBACH1^OE^*fish showed a greater proportion of Sox2^+^ cells at 7 dpi and fewer Sox2^+^ cells at 42 dpi **(Fig. 6G)**. These findings showed hBACH1 overexpression has an opposite phenotype to Bach1a/b loss-of-function, enhancing early activation and promoting late repression of Sox2^+^ cells after SCI. However, like *bach1a/b* mutants, *hBACH1^OE^*fish showed significantly lower swim endurance compared to controls **(Fig. 6H)**, suggesting tight regulation of *sox2* is critical for successful SC repair.

### *sox2* overexpression rescues SC regeneration in *bach1a/b* mutants

Our genetic and molecular studies indicated Bach1 regulates *sox2* expression and that *sox2* dysregulation in *bach1* mutants may underlie their regeneration defects. To test this hypothesis, we examined whether deficient regeneration in *bach1a/b* mutants could be rescued by *sox2* overexpression. To this end, we used an established *hsp70:sox2^x21^* transgenic line, referred to hereafter as *sox2^OE^*, to induce *sox2* expression under control of a heat shock inducible promoter (Millimaki et al., 2010) **(Fig. 6I, 6J)**. We crossed the *sox2^OE^*transgene into *bach1a/b* mutants to generate *sox2^OE^*;*bach1a/b^-/-^*fish. *bach1a/b^-/-^* and *sox2^OE^*;*bach1a/b^-/-^*fish were subjected to SC transections and daily heat shocks to induce transgene expression. Compared to *bach1a/b^-/-^* fish where functional recovery is impaired, swim endurance and axon regrowth were significantly higher in *sox2^OE^*;*bach1a/b^-/-^* fish at 42 dpi **(Fig. 6K, S6C)**. These findings indicated that exogenous *sox2* expression partially rescues the regeneration defects in *bach1a/b* mutants and support a model in which tightly controlled Bach1-mediated *sox2* expression is required for successful SC repair.

## DISCUSSION

Adult zebrafish offer a valuable platform to elucidate transcriptional mechanisms of innate SC repair in regenerative vertebrates. By generating a multidimensional map of SC progenitors in zebrafish, this study provides a much-needed resource to determine the molecular underpinnings of progenitor cell potency and to advance translational studies. Anatomically and molecularly, we found *sox2^+^* cells are compartmentalized into niches of ependymal and non-ependymal progenitors that exhibit specific lineage biases in homeostatic and lesioned SC tissues. Temporally, our results support a model in which a single transcription factor like Bach1 can either activate or repress *sox2* expression to fine tune progenitor cell expansion and quiescence during the time course of regeneration.

SC progenitors are thought to maintain elevated potency in adult zebrafish, but not in mammals (Horky et al., 2006; Meletis et al., 2008; Reimer et al., 2008; Barnabe-Heider et al., 2010; Karimi-Abdolrezaee et al., 2012; Su et al., 2014; Becker and Becker, 2015; Ribeiro et al., 2017; Paniagua-Torija et al., 2018; Shah et al., 2018; Klatt Shaw et al., 2021; Saraswathy et al., 2022; Xue et al., 2022; Zhou et al., 2023). Yet, our understanding of the identities, heterogeneity and potency of these cells remains limited. Consequently, spinal progenitors have been termed ependymal, ependymo-radial glial, glial or neural stem cells, and interchangeably labeled with markers such as *sox2, gfap*, *hey, nes,* and *foxj1a* (März et al., 2010; Kroehne et al., 2011; Ogai et al., 2014; Ribeiro et al., 2017; Than-Trong et al., 2018; Than-Trong et al., 2020). By genetic lineage tracing, we showed that *sox2*^+^ cells give rise to most regenerating cell types after SCI. Our single-cell analysis and *in vivo* validations revealed ependymal and non-ependymal niches of progenitor cells in both homeostatic and lesioned SC tissues. Further studies will delineate the lineage relationships between different domains of *sox2* expressing cells. Consistent with prior knowledge, markers such as *foxj1a* and *nes* are expressed in restricted progenitor domains comprising ependymal and neural stem cells, respectively (Ribeiro et al., 2017; Xue et al., 2022). On the other hand, *gfap* is more broadly expressed in SC progenitors, but also in non-progenitor glia and other cell types (Lam et al., 2009; Goldshmit et al., 2012; Johnson et al., 2016). Here, we used *sox2* to comprehensively characterize SC progenitors, but it is important to note that *sox2* is also expressed in additional cell types after SCI. As there is no single definitive marker that is both inclusive and specific to all SC progenitors, we propose designating cell populations by the markers they express (e.g. *sox2*^+^ cells), rather than relying on alternate nomenclature.

At the core of zebrafish regenerative capacity and this study is the ability of SC tissues to curtail regenerative processes and restore quiescence as SC regeneration concludes. Restricting the proliferation of progenitor cells is essential to reestablish homeostasis and prevent the detrimental consequences associated with sustained cellular proliferation. Yet, most tissue regeneration research has centered on promoting proliferative repair studies (Becker et al., 2004; Reimer et al., 2013; Barreiro-Iglesias et al., 2015; Hui et al., 2015; Mokalled et al., 2016; Ribeiro et al., 2017; Cavone et al., 2021; Klatt Shaw et al., 2021; Vandestadt et al., 2021; Saraswathy et al., 2022; Cigliola et al., 2023; Zhou et al., 2023; de Sena-Tomas et al., 2024), while the regulatory mechanisms that reinstate quiescence post-regeneration are largely unknown. Our findings support a model in which specific Bach1-Maf interactions can either activate or repress *sox2* expression and progenitor cell expansion, ostensibly in response to extracellular cues that relay the tissue’s regenerative state. Insights from salamander midbrain regeneration revealed a negative feedback loop from regenerating dopaminergic neurons to neurogenic progenitors to re-establish homeostasis following regenerative neurogenesis (Berg et al., 2011). In this model, dopamine signaling from regenerated neurons or administration of a dopamine receptor agonist are sufficient to limit progenitor cell proliferation (Berg et al., 2011). Intriguingly, in mammals, application of either dopamine receptor agonist or antagonist have been shown to promote neural stem cell proliferation, but downstream mechanisms remain to be elucidated (Kippin et al., 2005; Yang et al., 2008). Further studies into the extrinsic signals that regulate the expression and activity of diverse Bach1 and Maf transcription factors are needed to better understand mechanisms of initiation and resolution of SC regeneration.

Dual-function transcription factors enable context-dependent transcriptional regulation during tissue regeneration. Although Bach1 is best known for its canonical repressive functions, it is increasingly recognized for maintaining stem-like states through transcriptional activation (Sun et al., 2002; Fuse et al., 2015; Sato et al., 2020; Niu et al., 2021). Our findings suggest Bach1’s canonical repressive functions downregulate *sox2* at late stages of regeneration, but that cofactor availability can override this repressive role at earlier time points. Unlike pioneer factors like Oct4 and Sox2, which simultaneously activate and repress different sets of genes to control stem cell maintenance versus differentiation, our results suggest Bach1 can directly repress or derepress the same target gene at different stages of regeneration (Zhang et al., 2019). We propose differential binding to Maff versus Mafg as one mechanism that underlies Bach1 dual functions. However, as an epigenetic regulator, Bach1 may also act by recruiting chromatin remodeling complexes to regulate *sox2* expression (Guo et al., 2023). Further characterization of Bach1 binding partners as well as chromatin accessibility at different stages of regeneration are needed to better understand the dynamics of gene regulation during SC regeneration.

## ACKNOWLEDGMENTS

We thank V. Cavalli, H. Gabel, P. Williams, L. Sheets (Washington University) for discussion; S. Nandagopal, T. Tsai (Washington University), S. Megason (Harvard) for the HCR probe design script; B. Riley (Texas A&M) for zebrafish lines. We thank the Zebrafish Consortium Facility at Washington University for animal care, and the Genome Access Technology Center at Washington University for single-cell sequencing. This research was supported by grants from the NIH (R35 GM118179 to LSK, R01 AR070299 to ANJ, R01 NS113915 and R01 NS123708 to MHM).

## METHODS

### Zebrafish

Adult zebrafish of the Ekkwill, Tubingen and AB strains were maintained at the Washington University Zebrafish Consortium Facility. All animal experiments were performed in compliance with institutional animal protocols. Fish stocks were tracked and catalogued using VMS-Fish software (Muraleedharan Saraswathy and Mokalled, 2025). Adult male and female animals of ∼2 cm in length (4-6 months of age) were used. Experimental fish and control siblings of similar size and equal sex distribution were used for all experiments. SC transection surgeries and regeneration analyses were performed in a blinded manner and 2 to 4 independent experiments were performed using different clutches of animals. *bach1a^stl664^, bach1b^666^* and *sox2-2A-sfGFP^stl84^* were generated as previously described (Shin et al., 2014; Klatt Shaw and Mokalled, 2021). *Tg(hspl:sox2)^x21^* fish were generously provided by Dr. Bruce B. Riley (Texas A&M)(Millimaki et al., 2010).

### Generation of *hsp70:hBACH1-2A-EGFP* zebrafish

The following primers were used to amplify hBACH1 cDNA: ClaI forward primer 5’-ccATCGATatgtctctgagtgagaactcggtttt-3’ and ClaI reverse primer 5’-ccatcgat atgactgataaatgtactactgatgagt-3’. The genomic fragments were cloned into PCR2.1-TOPO vector, then subcloned into ClaI digested PCS2-hsp70-2A-EGFP plasmid. Clones were co-injected into one-cell stage wild-type embryos with I-SceI. A minimum of 3 founders were isolated and propagated. The transgenic line was established and designated *Tg(hsp70:hBACH1-2A-EGFP)^stl692^*. Animals were analyzed as hemizygotes.

### Generation of *sox2-2a-CreER^T2^* zebrafish

The DNA sequence for *CreER^T2^* was obtained from pME minprom_creER^T2^ (PMID 36975217) and assembled with the previously established *sox2-2a* targeting vector (M2) (Shin et al., 2014), by Gibson assembly. The *sox2-2a-CreER^T2^* targeting vector was linearized by NaeI. *sox2* TALEN RNAs were previously published (Shin et al., 2014) and synthesized with the SP6 mMessage mMachine Kit. Briefly, one-cell embryos were injected with *sox2* TALEN RNAs at 35 ng/μl each and linearized *sox2-2a-CreER^T2^*targeting vector at 10 ng/μl in 1x injection buffer (0.1 M KCL, 0.003% Phenol Red). Injected embryos were raised to adulthood and *sox2-2a-CreER^T2^* founders were identified by F1 progeny screen using PCR-based screen with *CreER^T2^* specific primers (cre_geno_F1 TGCAGGGAGAGGAGTTTGTGTGC and cre_geno_R1 TGTGGGAGAGGATGAGGAGGAGC). The obtained F1 progeny were further analyzed and confirmed that *2A-CreER^T2^* is precisely inserted by sanger sequencing. The knock-in line was established and designated *(sox2-2a-CreERT2)^stl374^.* Animals were analyzed as hemizygotes.

### Spinal cord transection and treatments

Zebrafish were anaesthetized in 0.2 g/l of MS-222 buffered to pH 7.0. Fine scissors were used to make a small incision that transects the spinal cord 4 mm caudal to the brainstem region. Complete transection was visually confirmed at the time of surgery. Injured animals were also assessed at 2 dpi to confirm loss of swim capacity post-surgery.

For EdU administration, zebrafish were anaesthetized and subjected to intraperitoneal EdU injections. 12.5 mM EdU (Sigma 900584) diluted in PBS was used. Daily injections were performed and paraformaldehyde-fixed cryosections were generated.

### Histology

Transverse cryosections of 16 µm thickness were generated from paraformaldehyde-fixed SC tissues. Tissue sections were imaged using a Zeiss AxioVision compound microscope or a Zeiss Axioscan.Z1 slide scanner for *in situ* hybridization, and a Zeiss LSM 800 confocal microscope or a Zeiss Axioscan.Z1 slide scanner for immunofluorescence.

For EdU staining, sections were incubated with freshly prepared EdU staining solution (100 mM Tris at pH 8.5, 1 mM CuSO4, 200 µM Fluorescent azide and 100 mM Ascorbic) for 30 min in the dark at room temperature.

For immunohistochemistry, tissue sections were hydrated in PBT then blocked with 5% goat serum for 1 h at room temperature. For nuclear antigens, sections were treated with 0.2% TritonX-100 for 5 min prior to the blocking step. Sections were incubated overnight at 4 °C with primary antibodies diluted in blocking agent, then washed in PBT, and treated for 1 h in secondary antibodies diluted in blocking agent at room temperature. Primary antibodies used in this study were: rabbit anti-Sox2 (GeneTex, 124477, 1:250), mouse anti-GFAP (ZIRC, ZRF1; 1:500), chicken anti-GFP (Aves Labs, GFP-1020; 1:1000) and rabbit anti-PCNA (GeneTex, GTX124496; 1:500). Secondary antibodies used in this study were: Alexa Fluor 488, Alexa Fluor 594, and Alexa 647 goat anti-rabbit or anti-mouse antibodies (Jackson ImmunoResearch, 1:200, 103-545-155, 111-605-003, 111-585-003, 115-605-003 and 115-585-003). Hoechst staining was performed according to the manufacturer’s recommendation (Thermo Fisher Scientific, H3570).

The HCR RNA *in situ* hybridization protocol was adapted from Molecular Instruments (Choi et al., 2018; Schwarzkopf et al., 2021). Briefly, tissue sections were hydrated and pre-treated either with 0.2% TritonX-100 for 5 min or boiled in Citrate Buffer (10 mM Citric Acid, 0.05% Tween-20, pH 6.0) for 10 min. For blocking, sections were incubated in Hybridization buffer (Molecular Instruments) for 1 h at 37 °C, then incubated in pre-warmed sets of DNA probes diluted to 0.0015 pmol/uL in Hybridization buffer for 48 h at 37 °C. Washes were then performed with Wash Buffer (Molecular Instruments) at 37 °C followed by 5x SSCT (3 M NaCl, 0.3 M Sodium Citrate, 0.1% Tween-20, pH 7.0). For signal amplification, sections were incubated in Amplification buffer (Molecular Instruments) for 1 h at room temperature. Prior to amplification, h1 and h2 hairpins were snap-cooled and diluted 1:50 in Amplification buffer. Amplification proceeded overnight at room temperature in the dark. Samples were then washed in 5x SSCT, 5x SSC and PBT before proceeding to immunohistochemistry. HCR RNA probes used in this study (*bach1a* and *bach1b*) were designed using a python script and ordered as 50 pmol opools from IDT **(Table S1)**.

RNAscope *in situ* hybridization for *bach1a* and *bach1b* was performed according to the manufacturer’ protocol (Advanced Cell Diagnostics, 322300).

### Histology quantification

Counting of Edu^+^, Sox2^+^ or PCNA^+^ cells were performed using a customized Fiji script (adapting ITCN: Image based Tool for counting nuclei; https://imagej.nih.gov/ij/plugins/itcn.html). The Fiji script incorporated user-defined inputs to define channels (including Hoechst) and to outline SC perimeters. To quantify nuclei, the following parameters were set in ITCN counter: width, 15; minimal distance, 7.5; threshold, 0.2. Once nuclei were identified, user-defined thresholds of individual cell markers were used to mask the image and identify nuclei located inside the masked regions. *xy* coordinates were extracted for each nucleus for cell counting. Raw counts and *xy* coordinates from Fiji were processed using a customized R script. Two markers were considered overlapping if they shared nuclei with same *xy* coordinates.

Colocalization analyses were performed using the JACoP plugin in Fiji (Bolte and CordeliÈRes, 2006). The polygon selection tool was used to outline SC perimeters.

HCR *in situ* hybridization signals were quantified using a customized Fiji script (Saraswathy et al., 2024). SC perimeters were defined and user-defined thresholds were set to create an inverted mask. Background noise was either manually corrected or defined by thresholding a dedicated, unstained background channel. Integrated density was quantified using the “Analyze Particles” command. To quantify RNA signals inside *sox2*^+^ cells, an inverted mask was created after thresholding *sox2*^+^ cells, then added to the final quantification mask. Fluorescence signals inside this newly generated mask were considered *sox2*^+^ cell specific.

For all statistical data, data points represent individual animals and N numbers are indicated. Statistical tests are reported in figure legends for each panel. Multiple comparisons were performed within groups and significant observations were reported.

### Isolation of spinal cord nuclei from adult zebrafish

Nuclear isolation from adult SC tissues was performed as previously described (Matson et al., 2018; Saraswathy et al., 2024). Briefly, 3 mm SC tissue sections flanking the lesion site were collected from 50 adult zebrafish at 7 and 14 dpi. Corresponding 3 mm tissue segments were collected from uninjured control siblings. For tissue lysis, the detergent mechanical lysis protocol described by Matson et al. was performed. SC tissues were homogenized at low setting for 15 s. Density gradient separation using sucrose solution was used to sediment the nuclei from the supernatant. Final nuclear lysates were resuspended using 100 µl of resuspension solution (1x PBS + 2% BSA + 0.2 U/µl RNase inhibitor). Hoechst staining was performed to assess the quality of isolated nuclei based on their shape. Samples, in which more than 70% of nuclei were scored as ‘healthy’, were submitted for single nuclear RNA-sequencing.

### Single nuclear RNA-sequencing

For snRNA-seq, 30 µl resuspension of isolated nuclei at a concentration of ∼1000 nuclei/µl was submitted to the Genome Technology Access Center in the McDonnell Genome Institute of Washington University. Two biological replicates were used for each time point (0, 7 and 14 dpi). cDNA was prepared after GEM generation and barcoding, followed by the GEM-RT reaction and bead cleanup steps. cDNA was amplified for 11−13 cycles then purified using SPRIselect beads. Purified cDNA samples were then run on a Bioanalyzer to determine cDNA concentration. GEX libraries were prepared as recommended by the 10x Genomics Chromium Single Cell 3’ Reagent Kits User Guide (v3.1 Chemistry Dual Index) with appropriate modifications to the PCR cycles based on the calculated cDNA concentration. For sample preparation on the 10x Genomics platform, the Chromium Next GEM Single Cell 3’ Kit v3.1, 16 rxns (PN-1000268), Chromium Next GEM Chip G Single Cell Kit, 48 rxns (PN-1000120), and Dual Index Kit TT Set A, 96 rxns (PN-1000215) were used. The concentration of each library was accurately determined through qPCR utilizing the KAPA library Quantification Kit according to the manufacturer’s protocol (KAPA Biosystems/Roche) to produce cluster counts appropriate for the Illumina NovaSeq6000 instrument. Normalized libraries were sequenced on a NovaSeq6000 S4 Flow Cell using the XP workflow and a 50 × 10 × 16 × 150 sequencing recipe according to manufacturer protocol. A median sequencing depth of 50,000 reads/cell was targeted for each Gene Expression Library. The Metadata, raw and processed data associated with this dataset is availale at the NCBI Gene Expression Omnibus.

### Aligning snRNA-seq reads

After sequencing, the Illumina output was processed using the CellRanger (v6.0.0) recommended pipeline to generate gene-barcode count matrices. A custom reference genome was made with the “cellranger mkref” command, using the fasta file of zebrafish reference genome GRCz11 constructed from the Ensemble genome build (https://useast.ensembl.org/Danio_rerio/Info/Index) and the sorted Gene Transfer Format file (v4.3.2) from the improved zebrafish transcriptome annotation(Lawson et al., 2020). Base call files for each sample from Illumina were demultiplexed into FASTQ reads. Then, the “cellranger count” pipeline was used to align sequencing reads in FASTQ files to the custom reference genome. Both exon and intron sequences were included during the alignment. The filtered gene-barcode count matrices generated by “cellranger count” was used for downstream analysis.

### Quality control

DecontX package (Yang et al., 2020) (celda v1.16.1) was used to remove droplets containing aberrant counts of ambient mRNA. The function “decontx” was used on the raw “RNA” counts to obtain “decontX” count data. Contamination score was assigned to each cell after running “decontx”. Cells with contamination scores greater than 0.75 were filtered out and only cells with contamination scores less than 0.75 were included for downstream analysis. Furthermore, decontX counts were used as default for pre-processing and normalization. After ambient RNA removal, DoubletFinder (McGinnis et al., 2019) (v2.0.3) was used to identify doublets formed from transcriptionally distinct cells. Optimal pK values were calculated from the outputs of function “paramsweep_v3”. Then, “doubletFinder_v3” function was used to predict doublet cells, where 50 principal components and pN value of 0.25 were given as input along with the previously calculated pK values (0.26). All the cells predicted as doublets were removed.

### Integrated analysis of snRNA-seq dataset

Datasets were integrated and analyzed using Seurat (v4.1.1) package with R (v4.2.1) (Team, 2018; Stuart et al., 2019). Each sample count matrix was filtered for genes that were expressed in at least 3 cells and cells expressing at least 200 genes, followed by cell quality assessment using commonly used QC matrices (Ilicic et al., 2016). Cells having a unique number of genes between 200 and 4000 and a mitochondrial gene percentage < 5% were used for downstream processing. Each dataset was independently normalized and scaled using the “SCTransform” function, which is an improved method for normalization, that performs a variance-stabilizing transformation using negative binomial regression (Hafemeister and Satija, 2019). Standard integration workflow of Seurat was used to identify shared sources of variation across experiments as well as mutual nearest neighbors(Butler et al., 2018; Haghverdi et al., 2018). Integration features were selected based on the top 4000 highly variable features using “SelectIntegrationFeatures” function (nfeatures = 4000), which was used as input for the “anchor.features” argument of the “FindIntegrationAnchors” function. PCA analysis was performed on the 4000 variable features and the top 50 principal components selected based on the elbow plot heuristic, which measures the contribution of variation in each component. These 50 principal components were used in “FindNeighbors” and “FindClusters” functions to perform graph-based clustering on a shared nearest neighbor graph(Levine et al., 2015; Xu and Su, 2015). Louvain algorithm was used for modularity optimization in cell clustering using “FindClusters” function. The resolution parameter (res = 0.4) that determines the granularity of clustering was selected by visually inspecting clusters with resolutions ranging between 0.1 and 2.0 as well as clustree graphs (Zappia and Oshlack, 2018). Uniform Manifold Approximation and Reduction (UMAP) was used for non-linear dimensional reduction of the first 50 principal components and visualize the data using “RunUMAP” function(Becht et al., 2018). Data was graphed using different plot functions, such as “DimPlot”, “VlnPlot”, “FeaturePlot”, “Dotplot” and “DoHeatmap”, to view the cell cluster identity and marker gene expression. Cell proportions were extracted using the “table” and “prop.table” functions. Differential gene expression for individual clusters was identified using Wilcoxon rank sum test in the “FindAllMarkers” function. Marker genes detected in at least 25% of the clustered cells and with a logFC threshold of 0.25 were selected. Only positive markers were reported.

### Subset of *sox2^+^* clusters

*sox2^+^* cells identified from the complete dataset were subclustered using the “subset” function for subcluster analysis. The *sox2^+^* subset was again normalized and scaled using the “SCTransform” function with glmGamPoi method (Ahlmann-Eltze and Huber, 2021). 50 principal components were used, and the resolution parameter was set to 0.7. Downstream analysis was done as described above for the integrated analysis.

### Cluster identification using differentially expressed markers

A “CNS markers” database of previously published markers of the different cell types that comprise the vertebrate brain and/or SC tissues was compiled(Saraswathy et al., 2024). For each cell cluster, every marker gene identified as a top differentially expressed (DE) marker of that cluster was cross-referenced with our compiled VNM database using our “DE-Marker-Scoring” algorithm (Saraswathy et al., 2024). For every matching marker gene, one point was given to the respective cluster under the column name with matching cell identity. Iteration over every marker gene was performed to generate a scoring matrix with varying points for each cluster against the different cell identities compiled in the VNM database. The “phyper” function in R was then used to calculate one-tailed binomial probabilities using hypergeometric distribution for the total score obtained from each cluster against each cell identity in the database. −log10 of probability values were obtained for plotting the heatmap. The resulting values were scaled from 0 to 100 and plotted as a heatmap using GraphPad prism. Each cluster was given an identity based on the maximum−log10 P score obtained in the heatmap. The top DE markers of clusters with ambiguous scores were manually searched in the literature (Zhang and Cui, 2014) (https://brainrnaseq.org/) to confirm cluster identity. “RenameIdents” function was used to assign identity to each cluster. To confirm the assigned cluster identities, enrichment of classical markers of respective cell types were tested using Dot plot.

### Selection of cluster markers

To identify gene expression markers for each *sox2^+^* cluster, top DE markers were filtered for specificity to such cluster. To this end, top DE markers were first filtered by their unique enrichment in one *sox2^+^* cluster **(Table S2, S3)**. Enrichment and percent expression of these unique genes were then confirmed using Dot plots. These top DE markers were ranked based on their specificity and cluster-specific expression enrichment. Then, each gene marker was visualized in the whole spinal cord dataset (including *sox2^-^* cells) using Dotplot and Featureplot to validate that they were indeed a specific marker for a *sox2^+^*cluster. At least two markers were selected for downstream HCR *in vivo* validation.

### Pseudobulk analysis

To identify enriched transcription factors, snRNA Seq data was aggregated using the “AggregateExpression” (Seurat v4.4) function for each replicate from uninjured, 7, 21 or 42 dpi time points (Saraswathy et al., 2024). Transcription factors were then filtered and classified by GO terms “TF.Activator” and “pos.regulation.tf” for transcriptional activators and “TF.Repressor” and “neg.regulation.tf” for transcriptional repressors **(Table S4)**. Transcription factors that are present in both classifications were categorized as “activator and repressor”. Differential expression analysis, compared to uninjured controls, was performed for each time point with DESeq2 using a p-value threshold of p<0.05 (Love et al., 2014). Only transcription factors with - 1>Log2(FC)>1 aggregate expression were considered down-or upregulated.

### Swim endurance assays

Zebrafish were exercised in groups of 8-12 in a 5 l swim tunnel device (Loligo, SW100605L, 120 V/60 Hz). After 10 min of acclimation inside the enclosed tunnel, water current velocity was increased every 2 min and fish swam against the current until they reached exhaustion. Exhausted animals were removed from the chamber without disturbing the remaining fish. Swim times at exhaustion were recorded for each fish. Results are expressed as mean ± SEM. One-way ANOVA and multiple comparisons were performed using the Prism software to determine statistical significance of swim times between groups.

### Axon tracing

Anterograde axon tracing was performed on adult fish at 42 dpi. Fish were anaesthetized using MS-222 and fine scissors were used to transect the cord 4 mm rostral to the lesion site. Biocytin-soaked Gelfoam Gelatin Sponge was applied at the new injury site (Gelfoam, Pfizer, 09-0315-08; biocytin, saturated solution, Sigma-Aldrich, B4261). Fish were euthanized 6 h post-treatment and biocytin was detected histologically using Alexa Fluor 594-conjugated streptavidin (Molecular Probes, S-11227). Biocytin-labeled axons were quantified using the ‘threshold’ and ‘particle analysis’ tools in the Fiji software. Four sections per fish at 600 µm (proximal) and 1500 µm (distal) caudal to the lesion core, and two sections 450 and 750 µm rostral to the lesion, were analyzed. Axon growth was normalized to the efficiency of biocytin labeling rostral to the lesion for each fish. Percentage axon growth was then normalized to the rostral level of the control group.

### Glial bridging

GFAP immunohistochemistry was performed on serial transverse sections from adult fish at 14 dpi. The cross-sectional area of the glial bridge (at the lesion site) and the area of the intact SC (750 µm rostral to the lesion) were measured using ImageJ software. Bridging was calculated as a ratio of these measurements.

### Quantitative RT-PCR (RT-qPCR)

RNA collections were performed using Trizol. 1 µg of RNA was converted into cDNA using the Maxima First Strand cDNA Synthesis Kit (Thermo Fisher Scientific, K1672) according to the manufacturer’s specifications. Quantitative PCR was performed using the Luna polymerase master mix (NEB, M3003) using gene-specific primers and the Bio-Rad CFX Connect Real-Time System **(Table S5)**. To determine primer efficiency, a standard curve was generated for each primer set using cDNA pooled from 7 dpi SC tissues. Only primer sets with a calculated amplification efficiency of 1.8-2.2 per cycle were used. Expression fold change for each gene was calculated using the ΔCq method and dual normalized to *eef1a1l1* gene expression and to target gene expression in control samples. For all RT-qPCR experiments, 2-3 biological replicates were used per genotype and a minimum of 3 SC tissues were pooled within each replicate. Each biological replicate was run in technical triplicate, and the average log_2_ fold change of these technical replicates is shown.

### Chromatin immunoprecipitation (ChIP)

*hsp70:hBACH1-2A-EGF=* zebrafish were subjected to SC transections and daily heat shocks for 1h at 33°C for 3 days prior to tissue collection. 4 mm SC tissues containing the lesion were collected at 5 or 42 dpi. ChIP was performed using the SimpleChIP Plus Sonication Chip Kit (Cell Signaling Technology #56383) according to the manufacturer’s instructions. Briefly, input SC tissues were cross-linked with 1% formaldehyde for 30 min, homogenized and sonicated. Chromatin fragment sizes were confirmed by gel electrophoresis. Immunoprecipitation was performed with either BACH1 polyclonal antibody (Proteintech #14018-1-AP) or rabbit IgG negative control antibody (Cell Signaling Technology #2729). Samples were incubated with primary antibodies overnight at 4°C, then with magnetic beads for 3h, washed and eluted. Candidate BACH1 binding sites were assessed by RT-qPCR using primers targeting candidate genomic regions containing the consensus BACH1 binding motif **(Table S5)**. Reported values represent BACH1 immunoprecipitated enrichment in *Tg^+^* animals normalized to Rabbit IgG control enrichment. A minimum of 12 SCs were pooled in each replicate.

### Luciferase Assay

Human Embryonic Kidney 293 (HEK293) cells (atcc #CRL-1573) were cultured in DMEM/F-12 supplemented with 10% fetal bovine serum, 1x L-Glutamine and 500 U/mL of penicillin-streptomycin. 15×10^3^ cells were seeded per well in a 96-well flat-bottom microplate (Corning #07200566) and allowed to adhere for 36 h at 37°C. Cells were transiently transfected with different combinations pGL3-sv40-Firefly luciferase constructs encoding 6XBACH1 or the *sox2* enhancer regions identified by ChIP. Cells were co-transfected with pcDNA3 constructs encoding zebrafish Bach1a, Bach1b, Mafga, Mafgb or Maff, along with a Renilla luciferase control. Cells were transfected using Lipofectamine-2000 following the manufacturer’s protocol (Invitrogen #15338100). Luciferase activity was evaluated using the Dual-Glo Luciferase Assay System (Promega #E2920) 36 hours after transfection. Firefly and Renilla Luciferase activity were measured using a Biotek Synergy HT plate reader. Reported values represent Firefly luciferase activity normalized to Renilla activity and pcDNA3 vector controls.

