## Supplemental Figures for "Transient activation of potent progenitor cells is required for spinal cord regeneration"

#### Figure S1

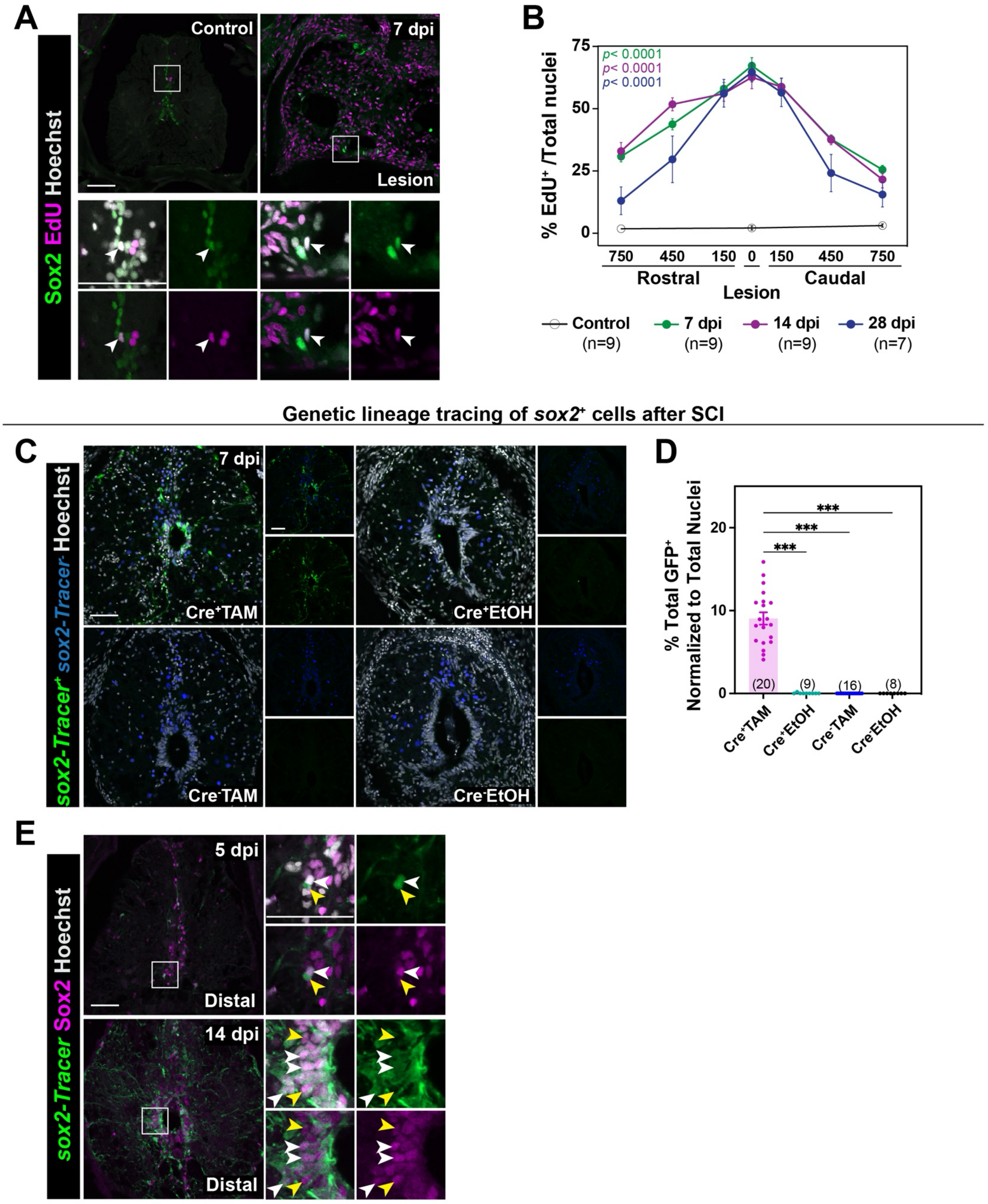
Proliferation of Sox2+ cells after SCI
A
Sox2 EdU Hoechst
Control
7 dpi
Lesion
B
% EdU+ /Total nuclei
p< 0.0001
p< 0.0001
p< 0.0001
75
50
25
0
750
450
150
0
150
450
750
Rostral
Caudal
Lesion
Control (n=9)
7 dpi (n=9)
14 dpi (n=9)
28 dpi (n=7)
Genetic lineage tracing of sox2+ cells after SCI
C
sox2-Tracer+ sox2-Tracer- Hoechst
7 dpi
Cre+TAM
Cre+EtOH
Cre-TAM
Cre-EtOH
D
% Total GFP+ Normalized to Total Nuclei
20
10
0
***
***
***
(20)
(9)
(16)
(8)
Cre+TAM
Cre+EtOH
Cre-TAM
Cre-EtOH
E
sox2-Tracer Sox2 Hoechst
5 dpi
Distal
14 dpi
Distal

**Figure S1. Proliferation and genetic lineage tracing of Sox2$^+$ cells during SC regeneration. (A)** Immunostaining for Sox2 (green), EdU (magenta) and Hoechst (grey) at 0, 7, 14 and 28 dpi. SC sections from uninjured control and 7 dpi animals are shown. Insets show high magnifications of Sox2$^+$ EdU$^+$ marked by white arrowheads. **(B)** Quantification of EdU$^+$ cells at 0, 7, 14 and 28 dpi. The numbers of EdU$^+$ cells were normalized to the total number of nuclei for each section. Cross-sections at 150, 450 and 750 µm rostral and caudal to the lesion were analyzed. **(C)** Immunostaining for GFP (*sox2*-Tracer$^+$ in green), mCherry (*sox2*-Tracer$^-$ in blue) and Hoechst (grey) at 7 dpi. SC sections from Tamoxifen-treated (TAM) *sox2*-Tracer (CreER$^{T2+}$) fish, TAM) CreER$^{T2-}$ fish, vehicle-treated (EtOH) CreER$^{T2+}$ fish and EtOH CreER$^{T2-}$ fish are shown. Single channel insets show *sox2*-Tracer$^+$ and *sox2*-Tracer$^-$ cells at a high magnification. **(D)** Quantification of *sox2*-Tracer$^+$ cells at 7 dpi. The numbers of *sox2*-Tracer$^+$ cells were normalized to the total number of nuclei for each section. Cross-sections 450 µm from the lesion were analyzed. **(E)** Immunostaining for *sox2*-Tracer$^+$ (GFP, green), Sox2 (magenta) and Hoechst (grey) at 5, 7, 14 and 28 dpi. SC sections from 5 dpi and 14 dpi animals are shown. Cross sections 450 µm from the lesion are shown. High magnification insets show select *sox2*-Tracer$^+$ Sox2$^+$ cells in triple, double or single channel views. White arrowheads indicate *sox2*-Tracer$^+$ Sox2$^+$ cells. Yellow arrowheads indicate *sox2*-Tracer$^-$ Sox2$^+$ cells or *sox2*-Tracer$^+$ Sox2$^-$ cells. Data points indicate individual animals and sample sizes are indicated in parentheses. Two-way ANOVA was performed in B. One-way ANOVA with Holm-Šidák's multiple comparisons were performed in D. Error bars represent SEM. ***p<0.001. Scale bars: 50 µm.

**Figure S2**

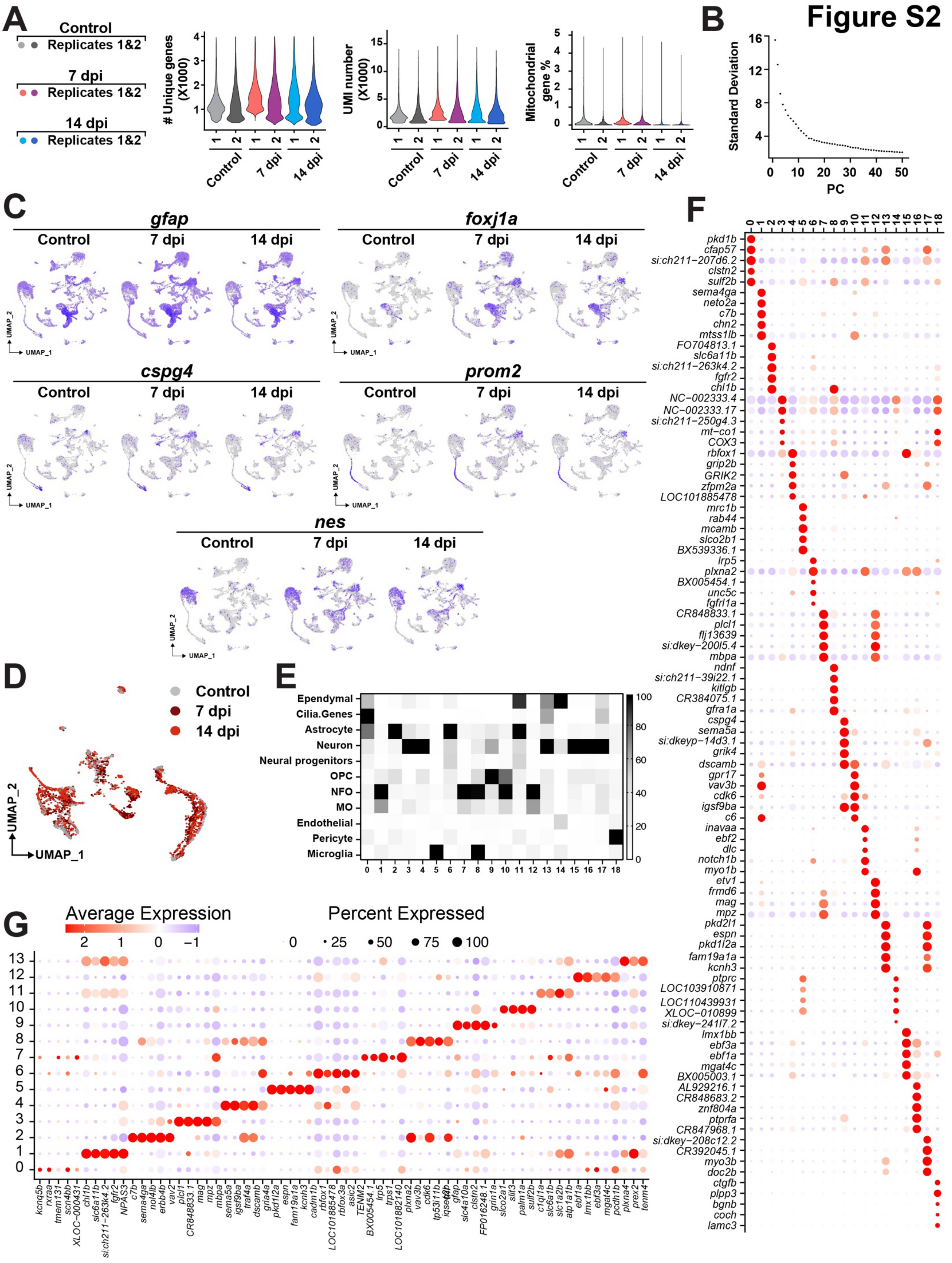

A
Control
Replicates 1&2
7 dpi
Replicates 1&2
14 dpi
Replicates 1&2
### Unique genes (X1000)
UMI number (X1000)
Mitochondrial gene %
Control
7 dpi
14 dpi
B
Standard Deviation
PC
C
gfap
foxj1a
cspg4
prom2
nes
Control
7 dpi
14 dpi
UMAP_1
UMAP_2
D
Control
7 dpi
14 dpi
UMAP_1
UMAP_2
E
Ependymal
Cilia.Genes
Astrocyte
Neuron
Neural progenitors
OPC
NFO
MO
Endothelial
Pericyte
Microglia
F
pkd1b
cfap57
si:ch211−207d6.2
clstn2
sulf2b
sema4ga
neto2a
c7b
chn2
mtss1lb
FO704813.1
slc6a11b
si:ch211−263k4.2
fgfr2
chl1b
NC−002333.4
NC−002333.17
si:ch211−250g4.3
mt−co1
COX3
rbfox1
grip2b
GRIK2
zfpm2a
LOC101885478
mrc1b
rab44
mcamb
slco2b1
BX539336.1
lrp5
plxna2
BX005454.1
unc5c
fgfrl1a
CR848833.1
plcl1
flj13639
si:dkey−200l5.4
mbpa
ndnf
si:ch211−39i22.1
kitlgb
CR384075.1
gfra1a
cspg4
sema5a
si:dkeyp−14d3.1
grik4
dscamb
gpr17
vav3b
cdk6
igsf9ba
c6
inavaa
ebf2
dlc
notch1b
myo1b
etv1
frmd6
mag
mpz
pkd2l1
espn
pkd1l2a
fam19a1a
kcnh3
ptprc
LOC103910871
LOC110439931
XLOC−010899
si:dkey−241l7.2
lmx1bb
ebf3a
ebf1a
mgat4c
BX005003.1
AL929216.1
CR848683.2
znf804a
ptprfa
CR847968.1
si:dkey−208c12.2
CR392045.1
myo3b
doc2b
ctgfb
plpp3
bgnb
coch
lamc3
G
Average Expression
Percent Expressed
kcnq5b
rxraa
tmem131
scn4bb
XLOC−000431
chl1b
slc6a11b
si:ch211−263k4.2
fgfr2
NPAS3
c7b
sema4ga
nol4lb
erbb4b
vav2
plcl1
CR848833.1
mag
mpz
mbpa
sema5a
igsf9ba
traf4a
dscamb
gria4a
pkd1l2a
espn
fam19a1a
kcnh3
cadm1b
rbfox1
LOC101885478
rbfox3a
asic2
TENM2
BX005454.1
lrp5
trps1
LOC101882140
plxna2
vav3b
cdk6
tp53i11b
iqsec3b
gfap
slc4a10a
clstn2
FP016248.1
grin1a
slco2a1
slit3
palm1a
sulf2b
c1ql1a
slc6a1b
slc1a2b
atp1a1b
ebf1a
lmx1bb
ebf3a
mgat4c
pcdh1b
plxna4
prex2
tenm4

**Figure S2. *sox2*[+] cells are heterogeneous and lineage-biased in homeostatic and lesioned SCs**. **(A)** Quality control metrics for snRNA-seq. Violin plots show the numbers of unique genes, numbers of unique molecular identifiers (UMI) and the proportions of UMIs that map to mitochondrial protein encoding genes. **(B)** Elbow plot shows ranking of principle components. **(C)** Feature plots showing the expression of select stem cell-related genes in the integrated dataset. **(D)** Merged UMAP representation of *sox2*$^+$ cells from 0, 7 and 14 dpi (color-coded by time point). **(E)** Lineage biases of *sox2*$^+$ cells in the integrated dataset. The top DE markers for each cluster were cross-referenced with our assembled database of vertebrate nervous system markers (VNM marker database). Heatmap depicting $-\log_{10}$ p-value for the marker scoring output from the "DEMarkerScoring" algorithm is shown. One-tailed hypergeometric probability test was used to calculate the p-value. $-\log_{10}$ p-value for each cluster was scaled between 0-100. Cluster identity was assigned based on the highest $-\log_{10}$ p-value obtained for each cluster. **(F)** Marker genes for subclusters of *sox2*$^+$ cells at 0, 7 and 14 dpi. Dotplot shows the top 5 markers for each subcluster. **(G)** Marker genes for subclusters of *sox2*$^+$ cells from uninjured SC tissues. Dotplot shows the top 5 markers for each subcluster. Dot colors and diameters represent average gene expression and percent cells with at least one UMI detected per gene, respectively.

Figure S3

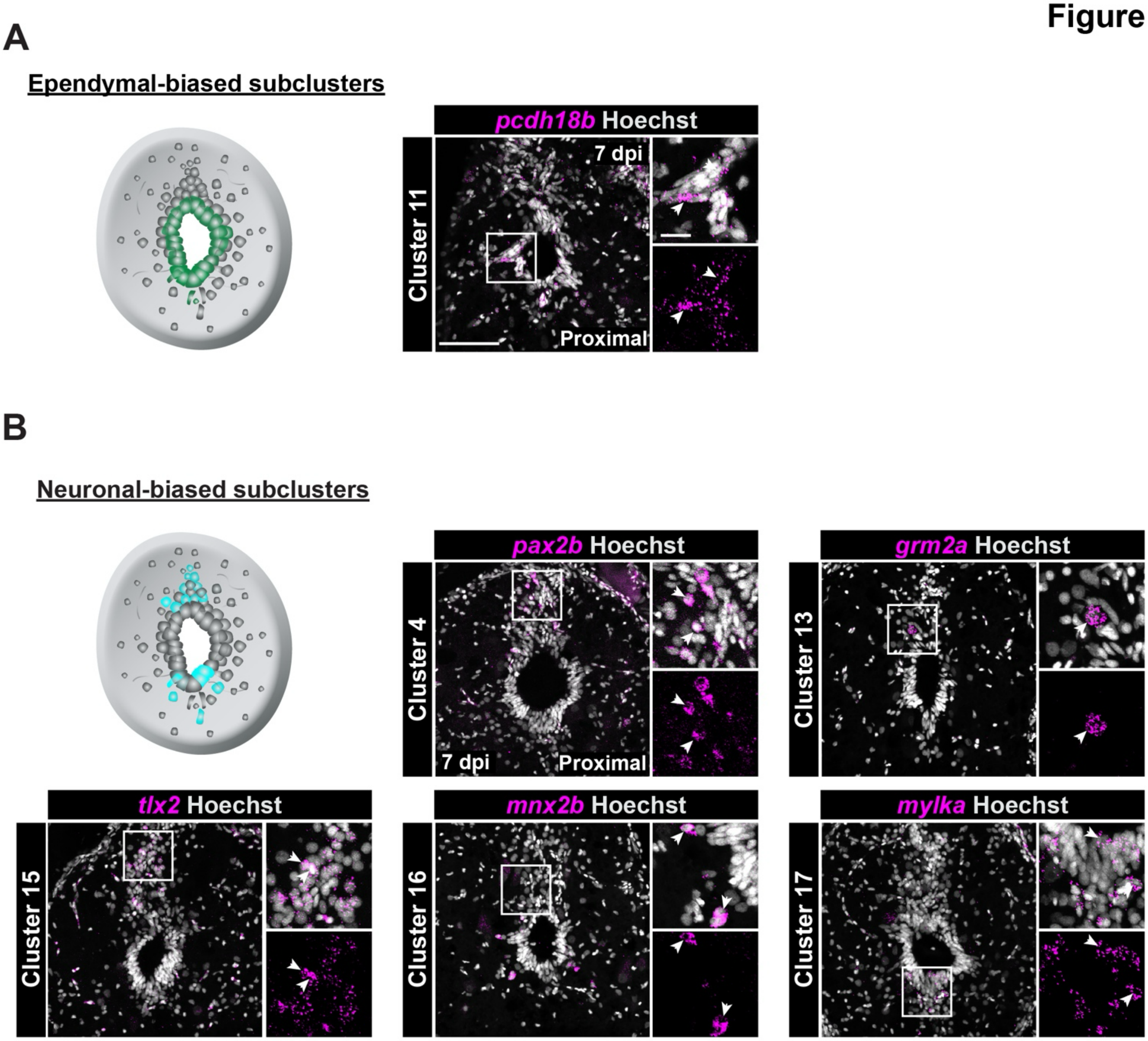

A
Ependymal-biased subclusters
pcdh18b Hoechst
Cluster 11
7 dpi
Proximal
B
Neuronal-biased subclusters
pax2b Hoechst
Cluster 4
7 dpi
Proximal
grm2a Hoechst
Cluster 13
tlx2 Hoechst
Cluster 15
mnx2b Hoechst
Cluster 16
mylka Hoechst
Cluster 17

**Figure S3. Mapping ependymal- and neuron-biased *sox2*$^+$ cells in adult zebrafish. (A)** *in vivo* validation of ependymal-biased *sox2*$^+$ cell cluster 11. HCR *in situ* hybridization for *pcdh18b* (magenta) and Hoechst (grey) at 7 dpi. Sections 450 µm from the lesion and high magnification insets are shown. **(B)** *in vivo* validation of neuron-biased *sox2*$^+$ cell clusters. HCR *in situ* hybridization for *pax2b* (cluster 4), *grm2a* (cluster 13), *tlx2* (cluster 15), *mnx2b* (cluster 16) and *mylka* (cluster 17) are shown. Sections 450 µm from the lesion and high magnification insets are shown. Arrowheads indicate positive cells. Scale bars: 50 µm, insets: 10µm.

### Figure S4

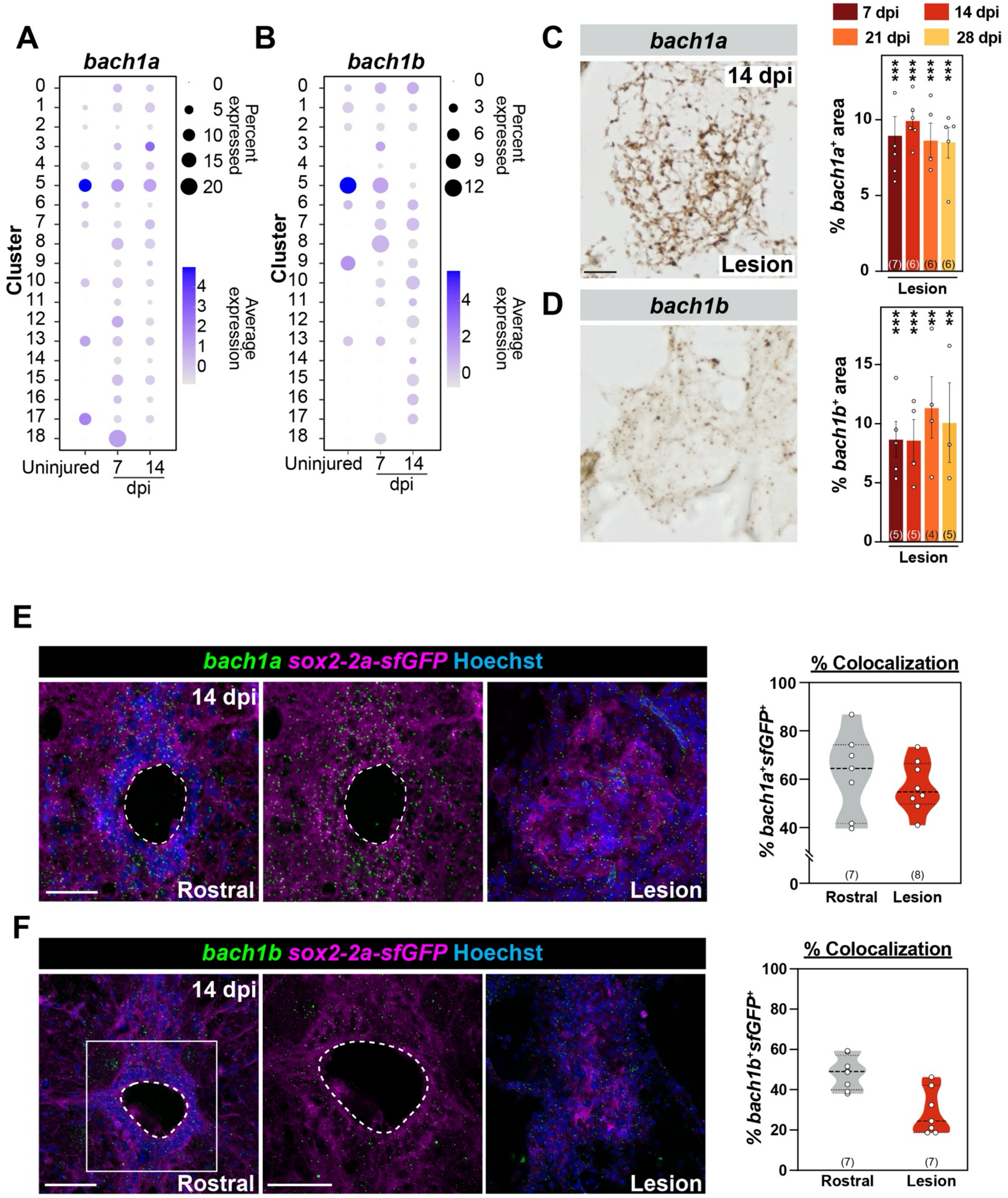

A
bach1a
Cluster
Percent expressed
0
5
10
15
20
Average expression
Uninjured
7
14
dpi
B
bach1b
Percent expressed
0
3
6
9
12
C
bach1a
14 dpi
Lesion
7 dpi
14 dpi
21 dpi
28 dpi
% bach1a⁺ area
D
bach1b
% bach1b⁺ area
E
bach1a sox2-2a-sfGFP Hoechst
14 dpi
Rostral
Lesion
% Colocalization
% bach1a⁺sfGFP⁺
(7)
(8)
F
bach1b sox2-2a-sfGFP Hoechst
% bach1b⁺sfGFP⁺

**Figure S4. *bach1a/b* expression during SC regeneration. (A,B)** Dotplot for *bach1a* and *bach1b* expression in each *sox2*$^{+}$ subcluster from Figure 2G at 0, 7 and 14 dpi. **(C,D)** RNAscope *in situ* hybridization for *bach1a* and *bach1b* during SC regeneration. SC sections at the lesion site are shown at 14 dpi. *bach1a* and *bach1b* expression at the lesion were quantified and normalized to uninjured controls. **(E,F)** HCR *in situ* hybridization for *bach1a* (E) and *bach1b* (F). SC cross sections 450 µm from the lesion are shown for *sox2-2A-sfGFP* zebrafish. Colocalization of *bach1a* or *bach1b* with *sox2-2A-sfGFP* at the lesion and 450 µm rostral to the lesion. For all quantifications, data points indicate individual animals and sample sizes are indicated in parentheses. Unpaired t tests were performed in C and D. Error bars represent SEM. **p<0.01; ***p<0.001. Scale bars: 50 µm.

### Figure S5

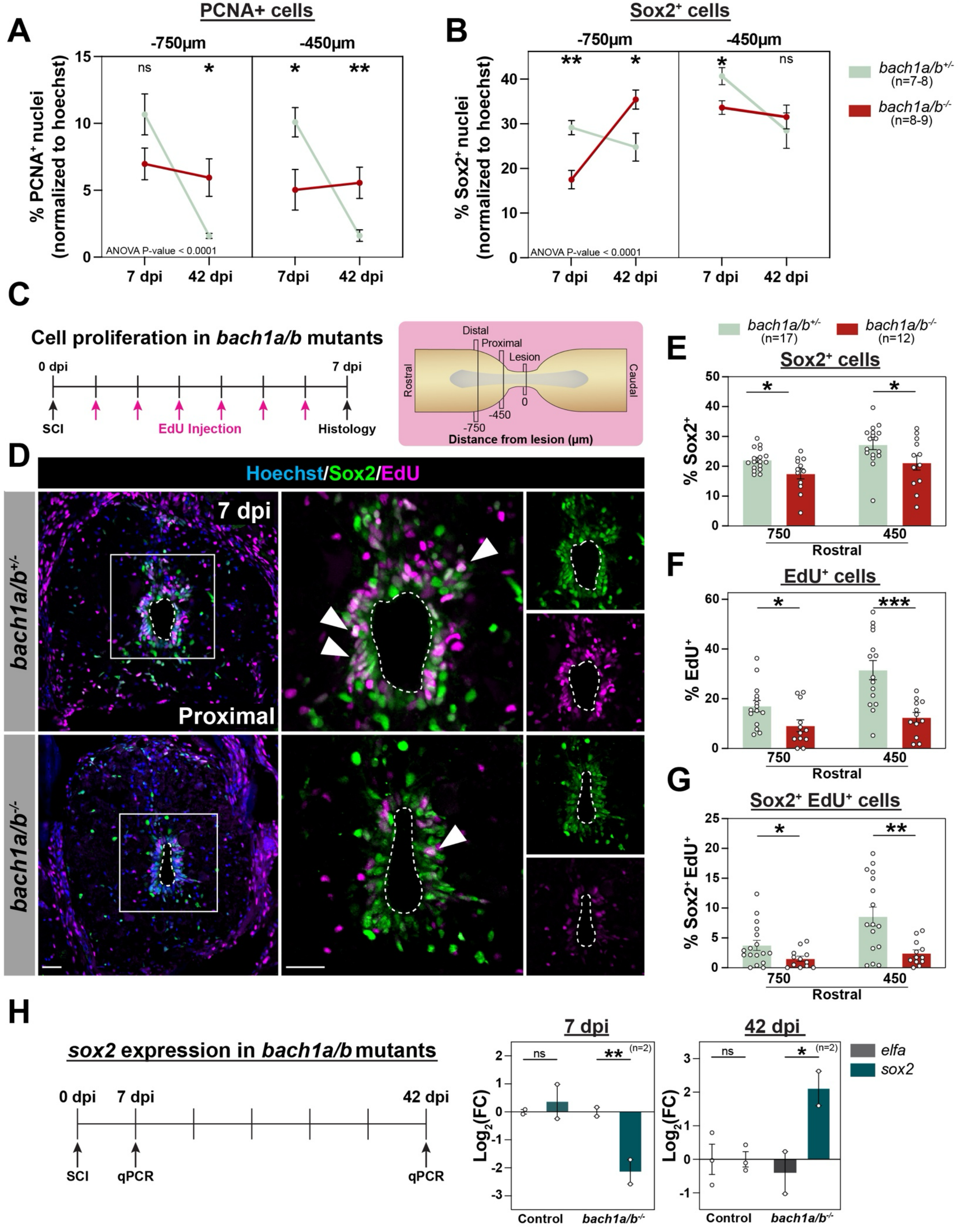

A
PCNA+ cells
-750µm
-450µm
% PCNA+ nuclei (normalized to hoechst)
ns
*
*
**
ANOVA P-value < 0.0001
7 dpi
42 dpi
7dpi
42 dpi
B
Sox2+ cells
-750µm
-450µm
% Sox2+ nuclei (normalized to hoechst)
**
*
*
ns
ANOVA P-value < 0.0001
7 dpi
42 dpi
7 dpi
42 dpi
bach1a/b+/- (n=7-8)
bach1a/b-/- (n=8-9)
C
Cell proliferation in bach1a/b mutants
0 dpi
7 dpi
SCI
EdU Injection
Histology
Distal
Proximal
Lesion
Rostral
Caudal
-750
-450
0
Distance from lesion (µm)
D
Hoechst/Sox2/EdU
7 dpi
Proximal
bach1a/b+/-
bach1a/b-/-
E
bach1a/b+/- (n=17)
bach1a/b-/- (n=12)
Sox2+ cells
% Sox2+
750
450
Rostral
F
EdU+ cells
% EdU+
*
***
750
450
Rostral
G
Sox2+ EdU+ cells
% Sox2+EdU+
*
**
750
450
Rostral
H
sox2 expression in bach1a/b mutants
0 dpi
7 dpi
42 dpi
SCI
qPCR
qPCR
7 dpi
42 dpi
Log2(FC)
ns
**
*
(n=2)
Control
bach1a/b-/-
elfa
sox2

**Figure S5. Loss of *bach1* impairs the expansion and quiescence of *sox2*$^+$ cells after SCI. (A,B)** Quantification of PCNA$^+$ cells (A) and Sox2$^+$ cells (B) in *bach1a/b* mutants at 7 and 42 dpi. For each section, the numbers of PCNA$^+$ cells or Sox2$^+$ cells were normalized to total nuclear counts. Quantifications were performed 450 µm from the lesion. **(C)** Experimental timeline to assess cumulative cell proliferation in *bach1a/b* mutants. Following SC transections, *bach1a/b*$^{-/-}$ and *bach1a/b*$^{+/-}$ control fish received daily intraperitoneal EdU injections. SC tissues were collected at 7 dpi for analysis. **(D)** EdU staining in *bach1a/b*$^{-/-}$ and *bach1a/b*$^{+/-}$ controls at 7 dpi. SC sections were co-stained for Sox2 (green) and Hoechst (blue). Representative micrographs 450 µm from the lesion are shown. Arrowheads indicate Sox2$^+$ PCNA$^+$ cells. **(E-G)** Quantification of Sox2$^+$, EdU$^+$ and Sox2$^+$EdU$^+$ cells in *bach1a/b*$^{-/-}$ and *bach1a/b*$^{+/-}$ controls. SC sections 450 and 750 µm from the lesion were quantified. For each section, the numbers of Sox2$^+$, EdU$^+$ or Sox2$^+$EdU$^+$ cells were normalized to total nuclear counts. Data points indicate individual animals and sample sizes are indicated in parentheses. **(H)** Experimental timeline to assess *sox2* expression in *bach1a/b*$^{-/-}$ and *bach1a/b*$^{+/-}$ controls. qRT-PCR assessed *sox2* expression in *bach1a/b*$^{-/-}$ and *bach1a/b*$^{+/-}$ control SCs at 7 and 42 dpi. Relative gene expression was calculated using the ΔΔCt method, following primer efficiency correction and normalization to *elfa*. Two-way ANOVA with Holm-Šidák's multiple comparisons were performed in A, B and E-H. Error bars represent SEM. ns, $p \geq 0.05$; *$p<0.05$; **$p<0.01$; ***$p<0.001$. Scale bars: 50 µm.

### Figure S6

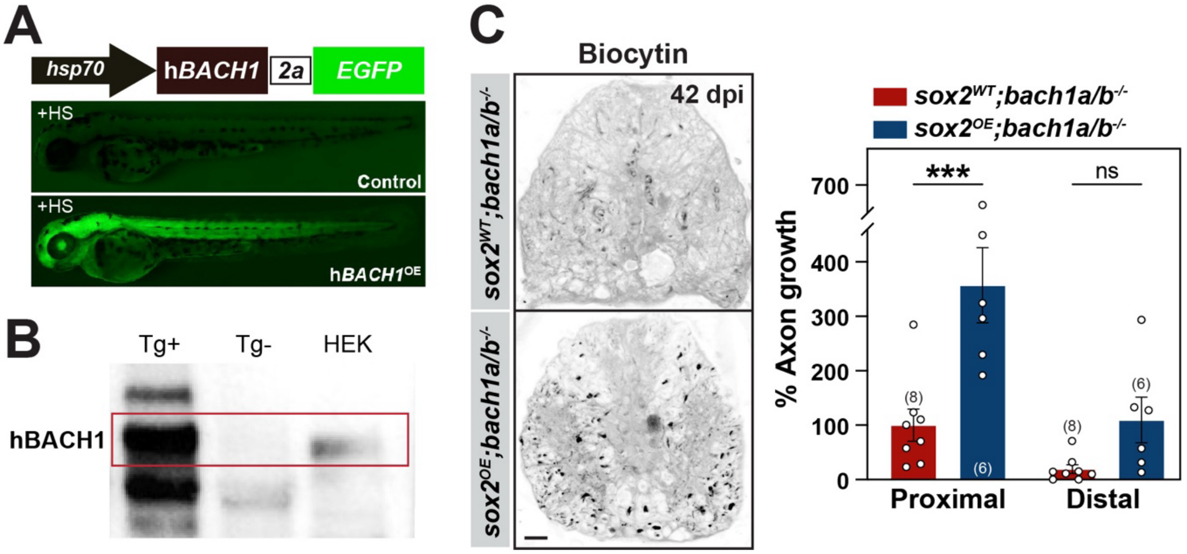
A
hsp70
hBACH1
2a
EGFP
+HS
Control
+HS
hBACH1OE
B
Tg+
Tg-
HEK
hBACH1
C
Biocytin
42 dpi
sox2WT;bach1a/b-/-
sox2OE;bach1a/b-/-
% Axon growth
700
400
300
200
100
0
***
ns
(8)
(6)
(8)
(6)
Proximal
Distal

**Figure S6. Bach1 is a dual regulator of *sox2* expression in sub-acute and chronic SCI. (A)** Generation of *hsp70:hBACH1-2A-EGFP* transgenic zebrafish. GFP expression is shown for *Tg*$^{+}$ and *Tg*$^{-}$ larvae 2 hours after a single heat shock. **(B)** Western blot for hBACH1 in *hsp70:hBACH1-2A-EGFP* larvae at 5 dpf. *Tg*$^{+}$ and *Tg*$^{-}$ larval homogenates were collected after a 1 hour heat shock. Homogenate from HEK-293T cells was included as a positive control. **(C)** Anterograde axon tracing in *sox2*$^{OE}$*;bach1a/b*$^{-/-}$ and *bach1a/b*$^{-/-}$ fish at 42 dpi. Biocytin axon tracer was applied rostrally and analyzed at 600 μm (proximal) and 1500 μm (distal) caudal to the lesion. Representative images show the extent of Biocytin labeling 600 μm (proximal) caudal to the lesion. Axon growth was first normalized to the extent of Biocytin rostral to the lesion, and then to labeling in *bach1a/b*$^{-/-}$ controls. For all quantifications, data points indicate individual animals and sample sizes are indicated in parentheses. Two-way ANOVA with Holm-Šidák's multiple comparisons were performed in C and D. Unpaired t test was performed in D. Error bars represent SEM. ns, $p \geq 0.05$; ***$p < 0.001$. Scale bars: 50 μm.
